# Senescence suppresses the integrated stress response and activates a stress-enhanced secretory phenotype

**DOI:** 10.1101/2023.04.12.536613

**Authors:** Matthew J. Payea, Showkat A. Dar, Carlos Anerillas, Jennifer L. Martindale, Cedric Belair, Rachel Munk, Sulochan Malla, Jinshui Fan, Yulan Piao, Xiaoling Yang, Abid Rehman, Nirad Banskota, Kotb Abdelmohsen, Myriam Gorospe, Manolis Maragkakis

**Affiliations:** Laboratory of Genetics and Genomics, National Institute on Aging, Intramural Research Program, National Institutes of Health, Baltimore, MD 21224, USA

## Abstract

Senescence is a state of indefinite cell cycle arrest associated with aging, cancer, and age-related diseases. Here, using label-based mass spectrometry, ribosome profiling and nanopore direct RNA sequencing, we explore the coordinated interaction of translational and transcriptional programs of human cellular senescence. We find that translational deregulation and a corresponding maladaptive integrated stress response (ISR) is a hallmark of senescence that desensitizes senescent cells to stress. We present evidence that senescent cells maintain high levels of eIF2α phosphorylation, typical of ISR activation, but translationally repress production of the stress response transcription factor 4 (ATF4) by ineffective bypass of the inhibitory upstream open reading frames. Surprisingly, ATF4 translation remains inhibited even after acute proteotoxic and amino acid starvation stressors, resulting in a highly diminished stress response. Furthermore, absent a response, stress augments the senescence secretory phenotype, thus intensifying a proinflammatory state that exacerbates disease. Our results reveal a novel mechanism that senescent cells exploit to evade an adaptive stress response and remain viable.

## Introduction

Cellular senescence is a state of virtually permanent cell cycle arrest that typically occurs in response to sublethal genomic damage. Senescence is expected to primarily guard against carcinogenesis by forcing cells with potentially tumorigenic mutations out of the cell cycle (Lowe et al., 2004; Rodier and Campisi, 2011), though other functions for senescence in development (Munoz-Espin et al., 2013) and wound healing (Demaria et al., 2014; Wilkinson and Hardman, 2022) have also been described. Senescent cells become detrimental to human health during aging where they accumulate, and are linked to several aging related diseases (Bussian et al., 2018; Rouault et al., 2021; Shorter et al., 2022; Tuttle et al., 2020).

In addition to permanent cell cycle arrest, senescent cells undergo several other stable phenotypic changes including an enlarged and flattened morphology, reduced mitochondrial function, persistent DNA-damage signaling, and a dramatic increase in protein secretion termed the senescence associated secretory phenotype (SASP) (Coppe et al., 2008; Gasek et al., 2021; Gorgoulis et al., 2019; Kuilman et al., 2010). The most unexpected phenotypes of senescent cells is their profound resistance to apoptosis through a joint downregulation in apoptotic proteins and upregulation in survival pathways (Baar et al., 2017; Marcotte et al., 2004; Zhu et al., 2015). However, the mechanisms by which senescent cells maintain function during increases in stress and macromolecular damage remain understudied and must be substantive since senescent cells can survive indefinitely.

Here we coupled Tandem-Mass Tag (TMT) mass spectrometry, ribosome sequencing, and full-length, direct RNA nanopore sequencing (dRNA-Seq) in senescent, quiescent, and cycling states to discover senescence-specific regulation of translation and the proteome that directly dictates cell survival and secretory programs. Our analysis found that human senescent cells possess a unique basal increase in ER stress accompanied by an increase in eIF2α phosphorylation, a critical marker of the integrated stress response (ISR), that regulates ternary complex availability for translation initiation. Surprisingly, we find that a global decrease in translating ribosomes that alters the demand for ternary complex decouples the expression of the downstream major ISR transcription factor ATF4 from eIF2α phosphorylation. This decoupling leads to ineffective bypass of an inhibitory upstream open reading frame (uORF) in *ATF4* mRNA.

Our results show that the senescence-associated resistance to ATF4 translation and ISR pathway activation further extends to acute proteotoxic, oxidative, and starvation stressors that all elicit a highly limited adaptive response. Instead, we find that stress induces an alternative transcriptional program that exacerbates the SASP through increases in both mRNA and secreted protein. This stress-induced enhancement of the SASP occurs after even transient stress and is sustained for days after the stress is removed. Our work reveals a new mechanism that defines the complexity of senescent cell homeostasis in the presence of stress driven by a highly depressed translational activity. We also find that stress induces a sustained increase in the inflammatory potential of senescent cells with substantial implications to aging and senescence-associated diseases.

## Results

### Cellular senescence is characterized by persistent ER stress signaling but post-transcriptional inhibition of ATF4 activation

To perform a direct and quantitative comparison of the senescent, cycling, and quiescent proteomes, we used TMT isobaric labeling and mass spectrometry using IMR-90 human lung fibroblasts. We chose a DNA damage-induced model of senescence for study, elicited by treating cells with the topoisomerase II inhibitor etoposide as previously described (Anerillas et al., 2022; Narita et al., 2011). Senescent cells (Sen) were compared to both early-passage cycling cells (Cyc) and contact-inhibited quiescent cells (Qui), to control for changes associated with general cell cycle arrest. We validated the senescence phenotype using a multi-marker workflow (Gorgoulis *et al*., 2019). As expected, senescent cells had increased expression of cell cycle inhibitors (p16/CDKN2A, p21/CDKN1A) and decreased expression of LMNB1 relative to cycling cells (**Fig. 1A**). Similarly, they had increased levels of SASP factors, *CDKN1A* and *CXCL8* mRNA (**Fig. 1B**). Senescent cells were also highly positive for senescence-associated β-galactosidase activity (SA-βGal) compared to cycling cells, further confirming the senescence phenotype (**Fig. 1C, Fig. S1A)**.

**Figure 1.**
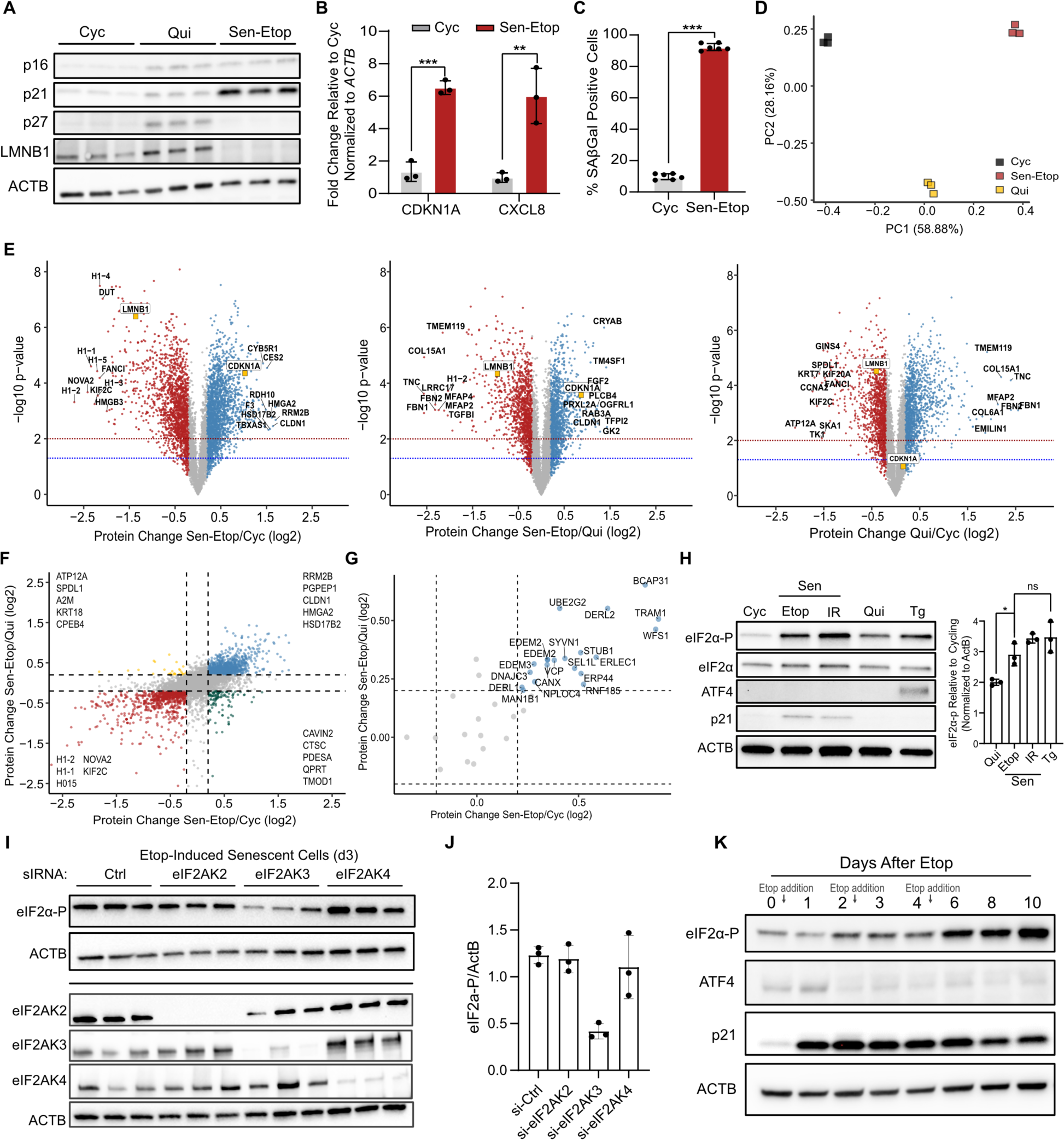
Cellular senescence is characterized by persistent ER stress signaling but no downstream ATF4 activation. **(A)** Western blot analysis of senescence markers p21 (CDKN1A), p16 (CDKN2A), and nuclear protein Lamin B1 (LMNB1) for etoposide induced senescent (Sen-Etop), quiescent (Qui), and cycling (Cyc) cells. ACTB is used as a loading control. **(B)** RT-qPCR analysis of *CDKN1A* and *CXCL8* mRNAs from cells analyzed in (A). Significance determined by t-test with n=3 (**= pvalue<0.01; *** = pvalue < 0.001) **(C)** Quantification of senescence-associated beta-galactosidase staining (SA-βGal) of Sen-Etop and Cyc cells (*** = pvalue < 0.001). **(D)** Principal component analysis of mass spectrometry data for cells harvested in triplicate. **(E)** Volcano plots of Sen-Etop/Cyc, Sen-Etop/Qui, and Qui/Cyc differential protein expression analysis, respectively. Senescence markers LMNB1 and CDKN1A colored in orange. Statistically significant (p-value ≤ 0.05 by two-tailed t-test) log2FC values are colored as follows: log2FC ≥ 0.2, blue; log2FC ≤ −0.2, red; log2FC between −0.2 and 0.2, grey. **(F)** Scatterplot of protein fold-change values from the Sen-Etop /Cyc (x-axis) and Sen-Etop /Qui (y-axis) comparisons. Dotted lines mark 0.2 and −0.2 log2FC in both comparisons. Quadrants are annotated with the top 5 most changed proteins, with respect to Sen/Cyc. **(G)** Scatterplot of protein fold-change values from the Sen-Etop /Cyc (x-axis) and Sen-Etop /Qui (y-axis) comparisons for selected ER stress protiens **(H)** Western blot analysis of ISR markers eIF2α phosphorylation (eIF2α-P) and ATF4; cells analyzed are cycling (Cyc), etoposide-induced senescent (Etop), ionizing-radiation induced senescent (IR), contact-inhibited quiescent (Qui), and cells treated for 3h with 25nM thapsigargin (Tg). At right, quantification of eIF2α-P signal relative to Cyc cells for Etop, IR, Qui, and Tg cells determined from western blot analysis in triplicate; normalized against ACTB protein expression. **(I)** Western blot analysis of senescent cells harvested 3d after transfection with siRNA targeting indicated eIF2α kinases (n=3). **(J)** Quantification of eIF2α-P signal normalized to ActB signal for western blot in **(I)**. **(K)** Western blot of lysates harvested at indicated time points after initial etoposide treatment. Etoposide treatments were applied at time points marked with arrows; harvests on day 0, 2, and 4 were performed prior to etoposide treatment.

Whole-cell protein lysates from the senescent, cycling, and quiescent cells were harvested in biological triplicate, processed for TMT-multiplex labeling and analyzed by LC/MS/MS. We identified 6,254 proteins across all 9 libraries **(Table S1)** and principal component analysis (PCA) showed that the proteomes of each cell type were highly distinct, whereas replicates were closely associated (**Fig. 1D**). We performed differential protein expression analysis for the senescent/cycling (Etop-Sen/Cyc), senescent/quiescent (Etop-Sen/Qui), and quiescent/cycling comparisons and defined a threshold for significant biological change that was retrospectively determined to be at least a ∼2-fold difference by western blot analysis **(Fig. S1B)**. As predicted by our PCA analysis, we identified several proteins with substantial change in all three comparisons (**Fig. 1E**). Senescent cells expressed 866 proteins that were uniquely decreased and 1,112 that were uniquely increased compared to both cycling and quiescent cells (**Fig. 1F, Table S1)**. Interestingly, gene ontology (Wu et al., 2021) and pathway enrichment analysis (Ge et al., 2020; Kanehisa et al., 2023) identified suppression of mRNA splicing and activation of protein processing in the endoplasmic reticulum (ER) as significant areas of senescence-specific change **(Fig. S1C-D)**. Indeed, our mass spectrometry data showed a clear senescence-specific increase in expression of ER stress associated proteins that included protein chaperones and other proteostasis components (**Fig. 1G**).

ER stress is monitored by the unfolded protein response (UPR) and the integrated stress response (ISR) pathway that co-activate to mediate an adaptive response to proteotoxic damage. We thus tested the activation of these pathways in two senescent models: etoposide-induced senescence (Etop-Sen), which was used above, and ionizing radiation-induced senescence (IR-Sen). As a positive control for ER stress, we treated cells with sub-lethal doses of the ER stress inducer thapsigargin (Tg) (Rutkowski et al., 2006). Expression levels of the UPR transcription factor XBP1 were not increased in senescent cells **(Fig. S1E)**, indicating that the UPR was likely not activated. However, analysis of the phosphorylation of eIF2α (eIF2α-P), an event on which ISR signals converge (Pakos-Zebrucka et al., 2016), revealed that senescent cells displayed high levels of eIF2α-P that was comparable to Tg-treated cells and significantly higher than in quiescent cells (**Fig. 1H**). To confirm the source of eIF2α-P we silenced the major ER stress kinase PERK (eIF2AK3). Silencing of PERK eliminated the increase in eIF2α-P, contrary to silencing GCN2 (eIF2AK4) and PKR (eIF2AK2), the kinases that primarily respond to ribosome collisions, amino-acid starvation, and viral infection (Pakos-Zebrucka *et al*., 2016; Wu et al., 2020) (**Fig. 1I-J**). Combined, our results show that senescent cells exhibit sufficient ER stress to induce eIF2α-P, a key step in the activation of the ISR.

Interestingly, while our results showed sufficiently high eIF2α-P levels in senescent cells, comparable to Tg-treated cells, we did not observe expression of ATF4, the core transcription factor of the ISR that is activated downstream of eIF2α-P (**Fig. 1H**). Given that the induction of senescence occurs in phases (Narita *et al*., 2011; Young et al., 2009) we tested whether ATF4 was expressed earlier during the establishment of senescence. While eIF2α-P levels rose with the induction of senescence (day 2) and remained elevated, ATF4 expression only briefly increased and then sharply declined during senescence establishment (**Fig. 1K).** This pattern of sustained eIF2α-P without ATF4 expression was consistent in IR-induced senescent cells, as well as in cells treated only once with etoposide **(Fig. S1F)**, suggesting a dynamic and progressive decoupling of ATF4 expression from eIF2α-P as senescence establishes. Since the senescence phenotype is highly heterogeneous with respect to both method of induction and cell type (Basisty et al., 2020; Casella et al., 2019; Hernandez-Segura et al., 2017) we examined whether increased eIF2α-P without ATF4 expression was generalizable to other models. We found that Etop treatment of WI-38 lung fibroblasts, BJ foreskin fibroblasts, and HUVEC vein endothelial cells as well as replicative exhaustion of IMR-90 cells also produced increased levels of eIF2α-P but not ATF4 **(Fig. S1G-H)**. Combined, our results show that senescence is generally associated with a persistent rise in ER-stress mediated eIF2α-P that is disconnected from downstream ATF4 production.

### Ribosome bypass of ATF4 uORF is not activated in senescent cells

Previous studies have shown that UV-induced DNA damage can increase eIF2α-P levels without increases in ATF4 through a reduction in *ATF4* mRNA levels mediated by the transcription factor C/EBPβ (Dey et al., 2010; Dey et al., 2012). We tested whether this could provide a mechanistic explanation for the lack of ATF4 expression in senescence. However, RT-qPCR analysis showed no statistically significant change in *ATF4* mRNA levels between cycling and senescent cells, albeit without the increase observed for Tg treated cells (**Fig. 2A**). Silencing C/EBPβ also produced no observable changes in *ATF4* mRNA or ATF4 levels **(Fig. S2A)**. Previous studies have established that the expression of the main ATF4 ORF is regulated by inhibitory uORFs that are only preferentially bypassed by ribosomes when the availability of ternary complex (eIF2α-GTP-tRNA_i_^Met^) for translation initiation is limited (Baird and Wek, 2012; Pakos-Zebrucka *et al*., 2016). Thus, efficient uORF bypass on ATF4 requires persistent ribosome demand accompanied by reduction of ternary complex supply, mediated by eIF2α-P. The disconnect between heightened eIF2α-P levels and ATF4 protein expression, despite stable *ATF4* mRNA abundance, raised the possibility that in senescence, the dynamics of this mechanism were altered.

**Figure 2.**
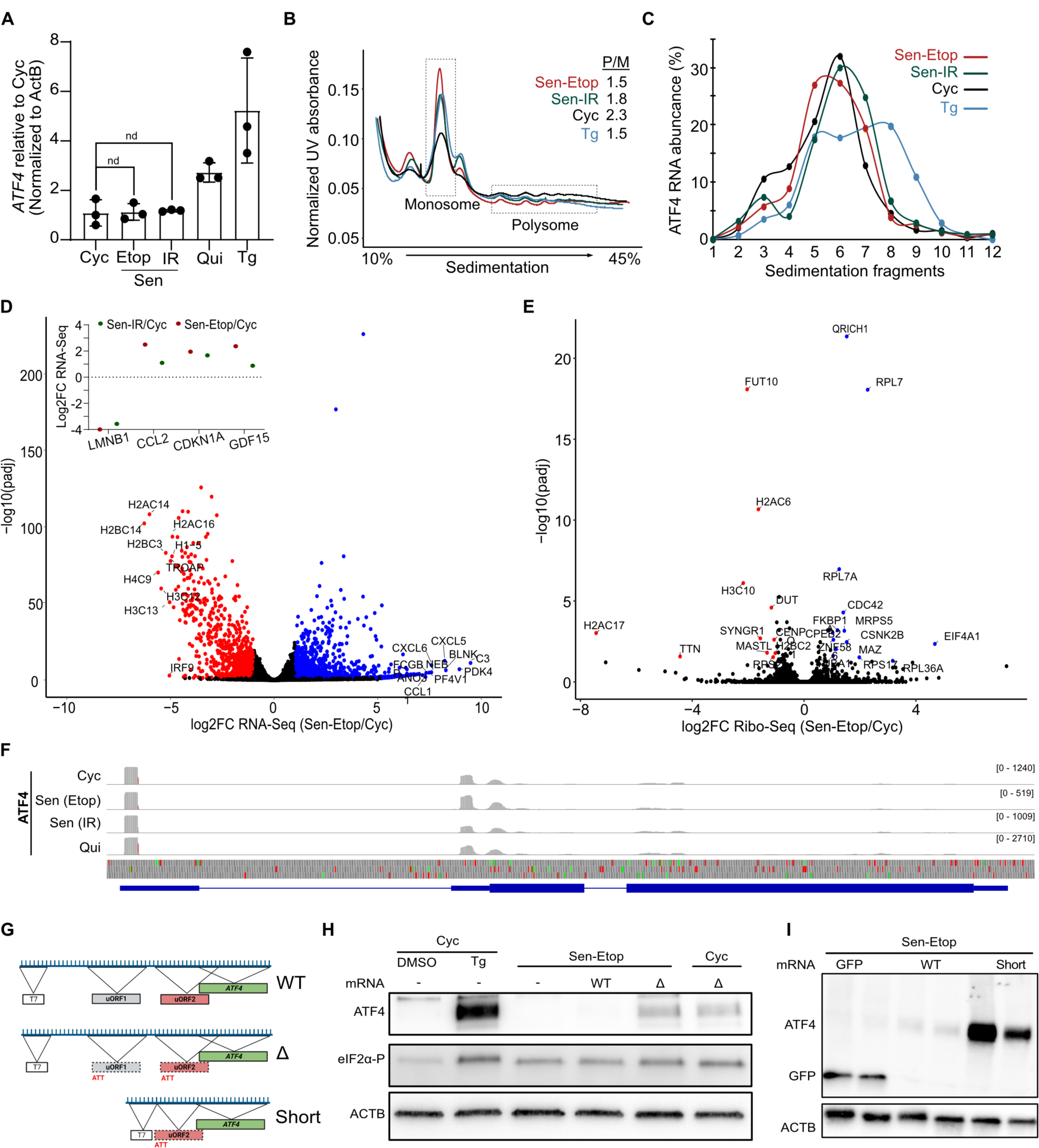
ATF4 translation is inhibited at uORF2 in senescent cells. **(A)** RT-qPCR analysis of *ATF4* mRNA levels. **(B)** Polysome profiling traces on a 10-45% sucrose gradient for lysates from indicated cell types after treatment with 0.1 mg/mL 22ycloheximide. The ratio of polysome/monosome fractions (P/M) are quantified and shown in inset. **(C)** RT-qPCR analysis of *ATF4* levels for RNA extracted from the polysome profiling experiment described in (B). **(D)** Differential RNA-seq expression analysis for the Sen-Etop/Cyc comparison. Statistically significant (adjusted p-value ≤ 0.05 by two-tailed t-test) log2FC values are colored as follows: log2FC ≤-1, red; log2FC ≥ 1, blue; log2FC between −1 and 1, black. Inset indicates log2FC values for senescence markers in the Sen-Etop/Cyc and Sen-IR/Cyc comparisons. **(E)** Volcano plot of Sen-Etop/Cyc differential translation efficiency (see methods) from ribosome sequencing. Statistically significant (adjusted p-value ≤ 0.05 by two-tailed t-test) log2FC values are colored as follows: log2FC ≤ −1, red; log2FC ≥ 1, blue; log2FC between −1 and 1, black. **(F)** Integrative Genome Viewer (IGV) mapping of ribosome footprints on the *ATF4* gene locus mapped using STAR (hg38 v.108). In translation track at bottom, green bars represent start codons and red bars represent stop codons. **(G)** Diagram of T7-*ATF4* mRNA constructs for transfection. T7 *in* vitro transcription site indicated as box, inhibitory uORF2 in red, and *ATF4* ORF in green; dotted lines indicate when uORF initiation sites have been mutated as indicated. **(H)** Western blot analysis of transfected *ATF4* mRNA for 6h as indicated for Cyc and etoposide-induced senescence (Sen-Etop). Cyc controls treated with DMSO or Tg (3h, 25 nM) in first two lanes. **(I)** Western blot of cells transfected with mRNA for 12h as indicated in etoposide-induced senescent (Sen-Etop) cells.

To test this hypothesis, we first performed polysome profiling for cycling, senescent, and Tg-treated cells normalized to total RNA amount. Our results showed that senescent cells had polysome distribution similar to Tg-treated cells with a characteristic decrease in translating polysomes (**Fig. 2B**). Interestingly treatment with the ISR inhibitory drug ISRIB that suppresses the effects of eIF2α-P showed reduced effect on senescent cells **(Fig. S2B-C)** compared to Tg cells, suggesting additional effects other than eIF2α-P for the reduction in translating ribosomes in senescence. To specifically probe the translational status of *ATF4* mRNA, we performed RT-qPCR analysis of each of the polysomal fractions. Our data showed that *ATF4* mRNA was predominantly in the non-translating monosome fraction of senescent and cycling cells, while it showed a clear shift toward heavy polysomes in Tg-treated cells (**Fig. 2C**). These results indicate that senescent cells do not engage *ATF4* mRNA in polysomes as would occur during eIF2α-P mediated uORF bypass.

To further test the translation dynamics on ATF4 and to globally assess why eIF2α-P may not function as expected in senescence, we performed ribosome sequencing (Ribo-Seq) of senescent, cycling, and quiescent cells, as described previously (McGlincy and Ingolia, 2017). Ribosome footprints (RPFs) were generated by incubating lysates with RNase I and the collapse of polysomes into monosomes was confirmed using polysome profiling **(Fig. S3A)**. RPFs were overwhelmingly (>85%) located within the coding sequence (CDS) of genes and had high periodicity **(Fig. S3B-C)** while replicates of Ribo-Seq and RNA-seq libraries, prepared in parallel, showed high correlation and PCA association **(Table S2, Fig. S3D, E)**. RNA-seq found several gene ontology terms ascribed to senescence **(Fig. S3F**) and showed that several previously described senescent markers (*CDKN1A*, *CCL2*, *GDF15, LMNB1*) displayed expression levels consistent with senescence (Casella *et al*., 2019; Hernandez-Segura *et al*., 2017) (**Fig. 2D**). Finally, it confirmed our initial finding that *ATF4* mRNA levels were not substantially changed in senescent relative to cycling cells **(Table S2).**

We next identified transcripts with detectable changes to translational efficiency relative to cycling cells. We found that both Etop and IR induced senescent cells had very few mRNAs (25 and 29, respectively) with a statistically significant (adjusted p-value ≤ 0.05) change in translation efficiency, though both were greater than what was observed in quiescent cells (5 mRNAs) (**Fig. 2E, Fig. S3G)**. Consistent with our western and polysome analysis, we found no significant difference in the translation status of *ATF4* mRNA in either of the four conditions **(Table S2)**. Interestingly, close examination of RPF distribution on *ATF4* mRNA showed that, in senescent cells, ribosomes were predominantly located in the uORFs rather than in the coding region (**Fig. 2F**) confirming that the expression of ATF4 remained suppressed by the inhibitory uORF2 in senescence despite increased eIF2α-P.

To confirm that ATF4 expression was repressed in senescence due to an impaired bypass and not due to insufficient mRNA levels, we overexpressed *in vitro*-transcribed *ATF4* mRNA constructs that were either unaltered (WT), possessed missense mutations in the start sites of uORF1 and uORF2 (ΔuORF), or were shortened up to uORF2 (Short) (**Fig. 2G**). We transfected the *in vitro* generated constructs into senescent and cycling cells and confirmed their overexpression using RT-qPCR analysis **(Fig. S3H)**. Transfection of senescent cells with the construct expressing WT *ATF4* mRNA produced no visible ATF4 signal, while transfection of the ΔuORF construct produced signals equivalent to those of similarly transfected cycling cells (**Fig. 2H**). Transfection with the Short construct resulted in an even higher increase in expression (**Fig. 2I**). Collectively, our results show that, even in the presence of elevated eIF2α-P and elevated *ATF4* mRNA, senescent cells are ineffective at bypass of the inhibitory uORF.

### Senescence is characterized by a global decrease in translating ribosomes

Our results suggested that resistance to ATF4 expression in senescence was due to a translational defect that prevented eIF2α-P from effectively mediating uORF bypass on ATF4. We searched our Ribo-Seq data for an aberration in the ribosome occupancy of the transcriptome of senescent cells using metagene analysis, but unexpectedly found that the distribution of ribosome footprints (RPFs) in the 5’ UTR, coding sequence (CDS), and 3’ UTR were nearly identical across all conditions (**Fig. 3A, Fig. S4A)**. Our polysome profiling distribution had suggested that senescent cells had fewer translating ribosomes (**Fig. 2B**). Due to normalization to RNA amount, it was unclear whether this was a result of a reduction in translating ribosomes or a shift from polysomes to monosomes. We therefore more closely examined the distribution of RPFs around the start site for all mRNAs to probe translation initiation. Interestingly, we saw that despite this shift, senescent cells had far fewer total RPFs than quiescent or cycling cells (**Fig. 3B, Fig. S4B)**, suggesting there were overall fewer translating ribosomes per mRNA in senescent cells.

**Figure 3.**
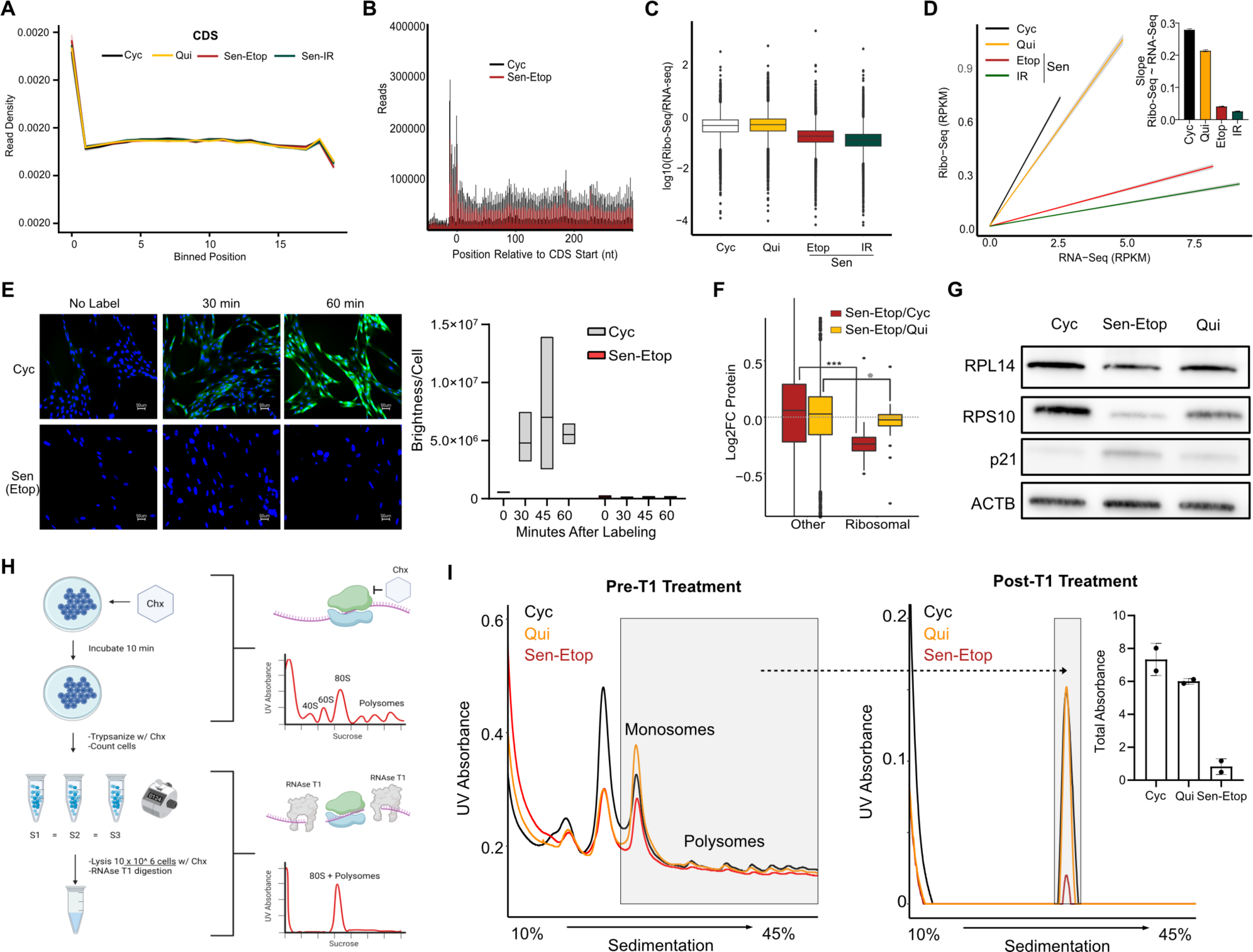
Senescent cells have a global decrease in translating ribosomes. **(A)** Metaplot analysis of ribosome footprints mapped to the transcriptome in the coding sequence (CDS) and 5’ and 3’ untranslated regions (5’ UTR and 3’UTR). Coordinates of footprints are aligned at the 5’ end and then binned for a transcriptome-wide metalength (see methods). **(B)** Metaplot distribution of ribosome footprints near the CDS start site (−100 to +300). Reads from etoposide-induced senescent cells (Sen-Etop) are compared to cycling cells (Cyc) and contact-induced quiescent cells (Qui). **(C)** Boxplot of log_10_(Ribo-seq/RNA-seq) reads as determined by RPKM analysis (see methods); **(D)**. Linear regression analysis of Ribo-seq∼RNA-seq RPKM reads with inset bar plot comparing slopes with standard error. **(E)** Representative images from protein synthesis pulse-label assay (see methods) where time indicates length of pulse. Green, nascent protein; blue, DAPI stain. At right, quantification of translation as brightness detected within nuclear region divided by total DAPI count detected (Brightness/Cell). **(F)** Boxplot of ribosomal protein abundances from mass spectrometry data. Significance of ribosomal protein reduction is assessed by student’s t-test (*= pvalue<0.01; *** = pvalue < 0.001). **(G)** Western blot validation of expression for select ribosomal proteins. **(H)** Diagram of ribosome content assay using T1 RNAse and cell count normalized lysates. **(I)** Polysome profiling using lysate derived from 10 million cells pre-T1 treatment (left) and post-T1 treatment (right); the grey boxes in both spectra are expected to represent the same total ribosomes. Post-T1 signal is quantified as the total absorbance within the grey box and summarized as a bar graph inset. **(J)** Model of the proposed continuum of ribosome and ternary complex availability that determines ISR activation in cycling, quiescence and senescence. The progressive imbalance of eIF2α/eIF2α-P ratio and ribosome abundance results in a persistent state of stress desensitization in senescence.

Indeed, further analysis showed that the ratio of Ribo-seq/RNA-seq reads normalized to total obtained reads revealed that senescent cells possessed a substantially depressed ribosome content (**Fig. 3C, Table S2)**. In fact, linear regression analysis of RNA-seq vs. Ribo-seq reads for the entire protein-coding transcriptome showed that senescent cells had ∼6 times less ribosomes per unit of RNA compared to both cycling or quiescent cells (**Fig. 3D**). As these results suggested that senescent cells might have markedly reduced total protein synthesis, we measured the rate of nascent protein production after pulsing with a methionine analogue. Both senescent models produced far less protein (green signal) over a 60-minute interval compared to cycling cells (**Fig. 3E, Fig. S4C)**. Combined these results suggested that senescent cells had overall fewer ribosomes.

While translational reductions are expected to accompany cell cycle arrest, the decreases we observed in senescence suggested they may exceed that of quiescent cells. To further explore these findings, we queried our mass spectrometry data for changes in components of the translation machinery. We observed that several translation-associated proteins were uniquely decreased in senescence **(Table S2)**, most striking among them were the ribosomal proteins that experienced a statistically significant (p-value ≤ 0.05) group-wide reduction in both the Sen-Etop/Cyc and Sen-Etop/Qui comparisons (**Fig. 3F**). The decrease in levels of ribosomal proteins was consistent with the observed reduction in ribosome footprints, as well as with broad decreases in rRNA biogenesis previously reported in senescence (Lessard et al., 2018; Nishimura et al., 2015). In fact, KEGG analysis of our mass spectrometry data revealed that senescent cells had an almost universal reduction of ribosome biogenesis components compared to both cycling and quiescent cells, particularly in proteins related to rRNA processing **(Fig. S4D)**. Western blot analysis confirmed decreases for 2 representative ribosomal proteins compared to cycling and cell cycle arrested quiescent cells (**Fig. 3G).**

To further test whether ribosome content in senescent cells was globally decreased, we modified our polysome profiling protocol to normalize to cell count, instead of RNA amount, and thus allow quantification of the number of ribosomes per cell. We subsequently treated lysates with T1 RNAse that degrades RNA but leaves ribosomes intact (**Fig. 3H**). Polysome profiling of lysates normalized to cell count before T1 treatment revealed a distinctly lower signal for senescent cells compared to quiescent and cycling cells that became quantifiable after treatment with RNAse T1 (**Fig. 3I).** Senescent cells showed an approximately 9-fold reduction in RNAse-resistant ribosomes compared to both cycling and quiescent cells (**Fig. 3I**), again showing that senescent cells had fewer ribosomes per cell than either cycling or quiescent cells.

The global reduction of ribosomes in senescent cell suggested a possible explanation for the ineffectiveness of eIF2α-P to induce ATF4 expression in senescence. Typically, eIF2α-P regulates expression of ATF4 by reducing concentration of the ternary complex and consequently increasing 40S scanning past the inhibitory uORF in ATF4. However, the global reduction in ribosomes we observe in senescent cells would be expected to demand less ternary complex for normal translation, thus effectively increasing the threshold of eIF2α-P required to induce ATF4 expression.

### Senescent cells fail to activate the ISR in response to acute stress

This model for ISR resistance predicts that further increasing eIF2α-P in senescence will have a diminished ability to increase ATF4 expression compared to quiescent and cycling cells. To test this, we subjected senescent, cycling, and quiescent cells to two distinct stressors known to increase eIF2α-P and activate the ISR: proteotoxic stress, through treatment with Tg as used above, and oxidative stress, through treatment with sodium arsenite (Ars). We found that cycling cells produced the strongest ATF4 expression, quiescent cells produced intermediate expression, and senescent cells produced minimal expression (**Fig. 4A**). We further quantified the efficiency of ISR-mediated activation of ATF4 by calculating the ATF4/eIF2α-P ratio of western blot signal, which confirmed that senescent cells consistently produced the least ATF4 expression in line with their highly reduced ribosome content (**Fig. 4B**). We next tested whether the reduced ATF4 expression of senescent cells was simply a change in the kinetics of ISR activation by analyzing a time course of Tg treatment. We saw that Tg treatment of cycling cells or cells rendered quiescent by serum starvation (SIQ) with Tg produced a rapid and sustained increase in eIF2α-P and ATF4, which was suppressed by addition of the eIF2AK3 inhibitor GSK2656157(Atkins et al., 2013) (**Fig. 4C).** Conversely, Tg treatment of IR and Etop induced senescent cells elicited only a minor increase expression of ATF4 that remained far below cycling and quiescent cells throughout the time course (**Fig. 4C, Fig. S5A)**. Quantification of the ATF4/eIF2α-P signal ratio again showed that senescent cells were ineffective at inducing ATF4 expression despite only modest differences in *ATF4* mRNA levels **(Fig. S5B)**. The senescence resistance to ATF4 expression further extended to treatment with the ER stress inducer tunicamycin **(Fig. S5B)**, where an even more extreme resistance was observed.

**Figure 4.**
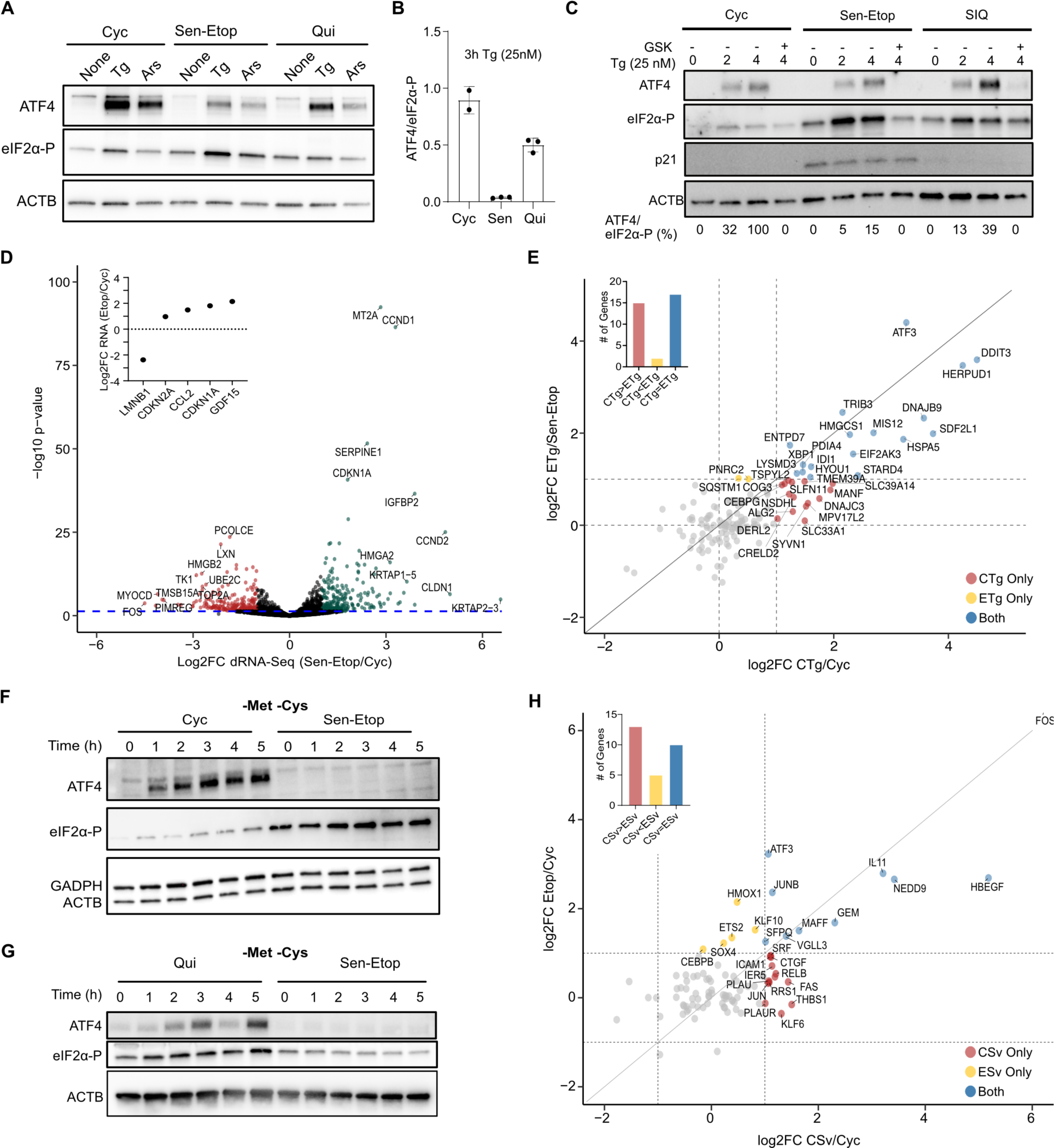
Senescent cells fail to activate the ISR in response to acute stress. **(A)** Western blot of etoposide-induced senescent (Sen-Etop), cycling (Cyc), and quiescent (Qui) cells treated for 3 h with DMSO (none), 25nM Tg (Tg), or 250 µM sodium arsenite (Ars). **(B)** The ATF4/eIF2α-P ratio is measured using intensity-based quantification of gel images and related as an integer percentage (right). **(C)** Western blot of Sen-Etop, Cyc, and serum-starved quiescent (SIQ) cells treated with 25 nM Tg and harvested after indicated times; eIF2AK3 inhibitor (GSK, 550 nM) added concurrently with Tg when indicated. The ATF4/eIF2α-P indicated at bottom, as described for (A). **(D)** Differential gene expression for nanopore-based direct RNA sequencing (dRNA-seq) of Sen-Etop and Cyc cells; green, log2FC ≥ 1, red, log2FC ≤ −1. **(E)** Scatter plot of differential gene expression for previously described ER-stress activated genes (Guan *et al*., 2017; Han *et al*., 2013; Shoulders *et al*., 2013). Dots are colored based on whether they are at least 2-fold increased (above dashed lines) in the indicated comparisons (exclusive or both). **(F-G)** Western blot of cells starved for indicated times in culture media lacking methionine and cysteine (-Met -Cys). **(H)** Scatter plot as in (D) for starvation activated genes (Tang *et al*., 2015; Tsalikis *et al*., 2016).

The lack of ATF4 expression in senescent cells, even in response to acute stress, suggested that the ISR as a whole was dysfunctional in senescence. To comprehensively test this possibility, we performed long-read direct RNA sequencing (dRNA-seq) (Ibrahim et al., 2021) of cycling and senescent cells with Tg treatment (CTg and ETg) compared to untreated cells (Cyc and Sen-Etop) **(Table S3, Fig. S5D)**. To our knowledge, there have been no previous analyses of senescent cells using a nanopore-based sequencing technology, and thus we first performed differential expression analysis between untreated senescent and cycling cells. Overall, we quantified 8,587 expressed genes **(Table S3, see methods)**, 271 of which showed statistically significant (adjusted p-value ≤ 0.05) mRNA abundance changes of at least 2-fold, including several senescent markers (Casella *et al*., 2019; Gorgoulis *et al*., 2019; Hernandez-Segura *et al*., 2017) (**Fig. 4D, Fig. S5E)**. Long-read sequencing also allowed us to interrogate features of the mRNAs that are not typically captured in short-read RNA-Seq analysis, such as differences in 5’ end degradation and poly-A tail deadenylation that may indicate changes in mRNA decay (Ibrahim *et al*., 2021). Our data showed little significant change (q-value ≤ 0.05) between cycling and senescent **(Fig. S5F)**. We next interrogated the expression of previously identified targets of the ISR (Guan et al., 2017; Han et al., 2013; Shoulders et al., 2013) in the Sen-Etop/Cyc and CTg/Cyc comparisons. As predicted, we found that while Tg treatment of cycling cells displayed 2-fold increases in several of these genes, only a few of them were elevated in Etop cells (**Fig. 4E**), strongly suggesting that the inability to elevate ATF4 production leads to a meaningfully decreased transcriptional response.

The ISR responds to numerous stressors, and therefore we tested whether senescent cells would also be resistant to a different form of ISR stress, triggered by amino acid starvation. We grew Cyc and Etop cells in media lacking the amino acids methionine and cysteine (-Met - Cys). We found that senescent cells produced no detectable ATF4 protein even after five hours of amino acid starvation, in contrast to Cyc cells, which responded within 1h (**Fig. 4F, Fig. S5G).** We further saw that the resistance of senescent cells to starvation again exceeded that of quiescent cells (**Fig. 4G**), supporting the view that reduced amino acid usage from cell cycle arrest was not the source of resistance by senescent cells. We again used dRNA-seq to analyze mRNA changes between Etop and Cyc libraries to cells starved of methionine and cysteine (ESv, CSv) **(Table S3)**. Evaluation of the starvation response by looking at mRNAs previously identified as being upregulated by starvation (Tang et al., 2015; Tsalikis et al., 2016) showed that ESv cells resulted in reduced expression compared to CSv (**Fig 4H**). Combined, our data strongly support our model for ATF4 resistance in senescence being a function of ribosome depletion and show that this resistance extends to a wholesale suppression of the ISR pathway even in the presence of acute exogenous stress.

### Stress remodels the senescence associated secretory phenotype

While senescent cells displayed a highly limited adaptive response to stress, we did observe senescence-specific increases in several transcripts, many of which encode proteins implicated in inflammatory pathways (**Fig. 5A**). The senescence associated secretory phenotype (SASP) is a hallmark trait of senescence whereby senescent cells secrete cytokines, interleukins, and other proteins (Basisty *et al*., 2020; Coppe et al., 2010; Coppe *et al*., 2008). To identify stress-induced increases comprehensively, we identified genes from our dRNA-seq data that were elevated after stress in senescent cells but not in cycling cells **(Table S3).** Pathway analysis identified terms associated with inflammatory and secretory pathways (**Fig. 5B**) similar to previous reports of ER stress association with an increase in inflammatory proteins (Kim and Lee, 2021; Lerner et al., 2012; Santarelli et al., 2020). We further refined our gene list to those annotated in the predicted secretome from the protein atlas (Uhlen et al., 2015). We found 21 secretory proteins, including several well-established SASP factors, among those encoded by stress-enhanced genes (**Fig. 5C**).

**Figure 5.**
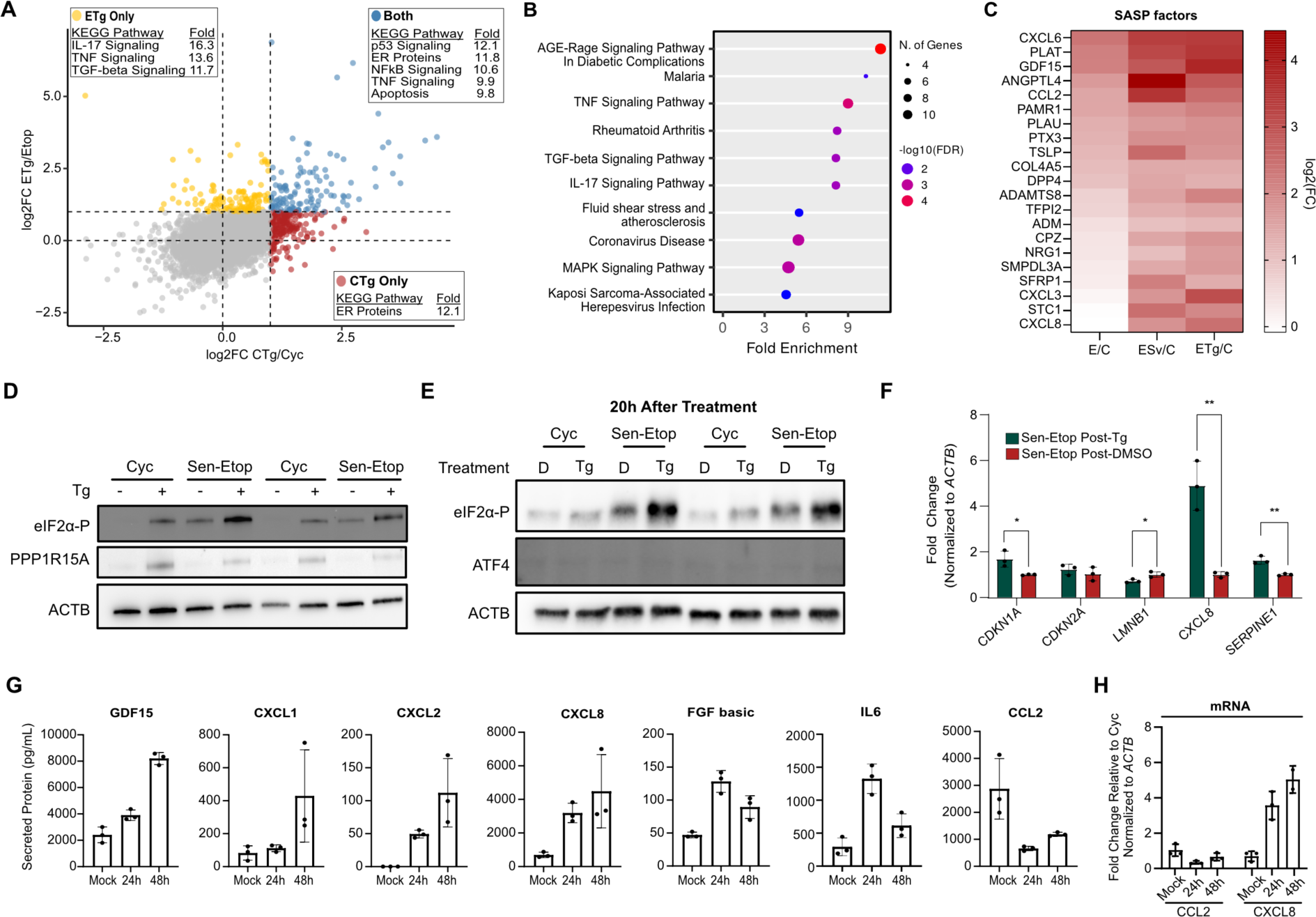
Acute stress remodels the senescence associated secretory phenotype. **(A)** Comparison of log2FC values for the ETg/Etop and CTg/Cyc comparisons for all detected RNAs; insets show enriched non-disease KEGG pathways for genes in respective quadrant. **(B)** KEGG pathway enrichment analysis of genes observed to be uniquely enhanced by stress(Kanehisa *et al*., 2023). **(C)** Heatmap of genes with annotated secretory function; color indicates log2FC for each library relative to cycling cells. **(D)** Western blot analysis of cells treated with or without 25nM thapsigargin for 3h (Tg). **(E)** Western blot of cells treated with DMSO (D) or Tg (as in (D)) and then allowed to recover for 20h in fresh media before harvest. **(F)** RT-qPCR analysis of cells treated in (E), significance assessed by student’s t-test (n=3, *= pvalue<0.01; ** = pvalue < 0.01). **(G)** Conditioned media collected in 24h intervals for cells treated with DMSO (Mock) or Tg for 3h at 25nM. Cells were then allowed to recover in fresh media for indicated time. Secreted proteins determined by a multiplex ELISA (see methods). **(H)** RT-qPCR of SASP factors *CCL2* and *CXCL8* from RNA extracted from cells used in (G).

In normal cells, activation of the ISR is followed by a recovery period after the initial stress has been resolved. This recovery process involves the action of the phosphatase GADD34 (PPP1R15A) that dephosphorylates eIF2α to enable the translation of the ATF4-activated transcripts (Pakos-Zebrucka *et al*., 2016). Western blot analysis showed that senescent cells expressed far less PPP1R15A than cycling cells after Tg treatment (**Fig. 5D**), suggesting that senescent cells may be deficient in the recovery from stress just as they are deficient in the activation of the adaptive response. In fact, we saw that even 20 h after Tg treatment, senescent cells maintained highly elevated levels of eIF2α-P, while cycling cells had mostly returned to a basal state (**Fig. 5E**). Interestingly, RT-qPCR analysis of these cells showed that the stress-induced SASP factors and senescent markers were also maintained after the stress was removed (**Fig. 5F**). To further examine this finding, we measured secreted proteins in the conditioned media of senescent cells allowed to recover after Tg treatment. Remarkably, we found that senescent cells maintained an elevated level of secretion up to even 48h following stress in a SASP-factor specific manner (**Fig. 5G**). The SASP is primarily regulated at the transcriptomic level (Coppe *et al*., 2008) and RT-qPCR analysis confirmed that changes in secreted protein levels were primarily consistent with underlying mRNA levels (**Fig. 5H**). Collectively our results show that acute stress induces a remodeling of the SASP that persists even after stress is withdrawn.

## Discussion

Senescence has been identified to serve multiple functions in organisms, from development to wound healing (Gorgoulis *et al*., 2019), but is primarily expected to serve as a guard against tumorigenesis induced through sub-lethal genomic damage. As such, senescence can be viewed as a stress response, since it detects genotoxic stress and then induces transcriptomic and proteomic remodeling for adaptation (Kuilman *et al*., 2010; Narita *et al*., 2011; Young *et al*., 2009). The principal component analysis of our dRNA-seq libraries point toward this same conclusion, wherein the first principal component appears to describe stress and places senescence as a more stressful state than exposure to either metabolic or proteotoxic stress (**Fig. 6A**). The substantial secretory protein synthesis of the SASP has previously been suggested as an additional source of stress in the ER of senescent cells (Dorr et al., 2013) Here we find that senescent cells indeed exhibit basal levels of ER stress that is sufficient to induce eIF2α phosphorylation, a converging step for ISR activation. The continuous nature of this stress suggests similarities to the chronic ISR where sustained ER stress in cycling cells results in changes to translation (Guan *et al*., 2017). However, an important distinction of senescence is that, unlike conventional stress responses that attempt to restore homeostasis or else induce apoptosis, senescence establishes a new and virtually permanent cell state that is in fact resistant to cell death. Thus, senescence has an entirely different set of demands from most other cell states: indefinite survival with unresolved cellular damage.

**Figure 6.**
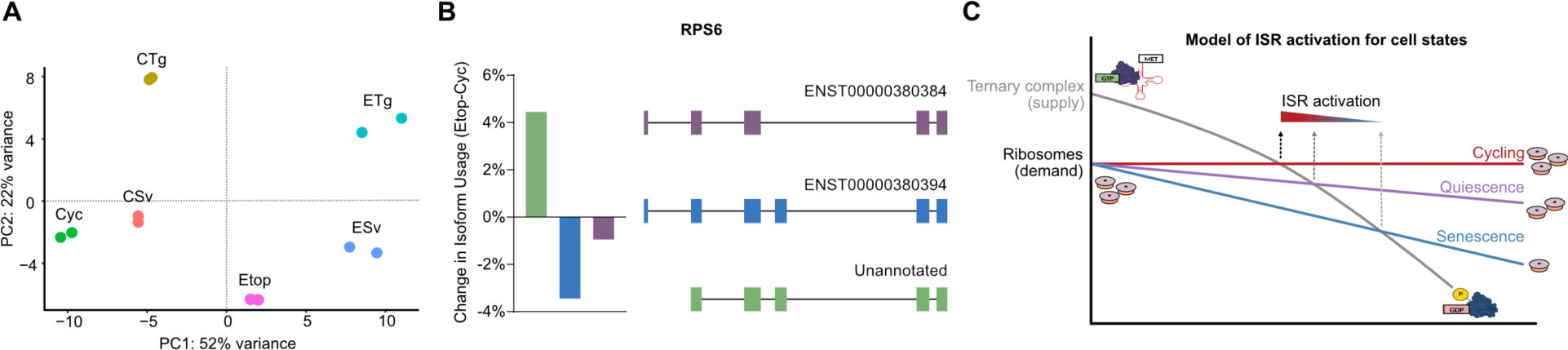
Model for translation repression and ISR inhibition in senescence. **(A)** Scatter plot of PC1 and PC2 from PCA of dRNA-Seq libraries. **(B)** Barplot of percent change in isoform usage and schematic of isoform structure for RPS6. **(C)** Model of the proposed continuum of ribosome and ternary complex availability that determines ISR activation in cycling, quiescence and senescence. The progressive imbalance of eIF2α/eIF2α-P ratio and ribosome abundance results in a persistent state of stress desensitization in senescence.

Part of this demand would involve a decline in translation, as the synthesis of proteins and biogenesis of ribosomes is the most energy intensive process of the cell (Proud, 2002; Schmidt, 1999). While reductions in translation are not novel to senescence, the extent to which senescent cells repress translation appears to exceed even that of quiescent cells and involves depletion of the ribosomes themselves. The mechanism by which this depletion is accomplished is not presently clear. Senescent cells expand in size which may dilute translational processes (Neurohr et al., 2019) and senescence has also been associated with decreases in rRNA transcription and maturation (Lessard *et al*., 2018; Nishimura *et al*., 2015). Interestingly, in addition to decreases in ribosomal protein production, our long-read sequencing data showed that ribosomal proteins frequently had increases in alternative splicing in senescence whose potential effects on ribosomes biogenesis still need to be explored (**Fig. 6B).** Alternatively, it is also possible that senescent cells accumulate inactive ribosomes, which are stable but not bound to mRNA, as have been recently shown to arise in several adverse cellular conditions (Gemmer et al., 2023; Smith et al., 2021).

While reduced translation and ribosome content has a predictable benefit to the conservation of cellular energy, we found it also has an unexpected effect on the activation of the ISR pathway. Our data show that the availability of translating ribosomes is a key determinant of senescent cell resistance to the ISR pathway, and even confers a moderate reduction in ISR sensitivity to cell cycle arrested quiescent cells This suggests a model of ISR activation that occurs in varying intensities over a continuum of ribosome content (**Fig. 6C**). The link between ribosome content and the efficacy of stress responses poses possible far-reaching implications to other conditions where ribosomes are dramatically decreased, such as ribosomopathies like Diamond-Blackfan anemia (Da Costa et al., 2021).

Stress also induced an unexpected change in the expression of inflammatory proteins in senescent cells even though they presented a limited adaptive response. This increase was sustained days after the initial stressor and was present at both the mRNA and secretome level. While these changes were often increases in protein secretion, there was high factor-specific variability, even among proteins that are typically closely associated, which suggests a more complex transcriptional program in response to stress. We call this phenomenon stress-induced secretory remodeling (SISR) to encompass the broad range of changes we observed. While previous studies have observed that hypoxic stress can mitigate the SASP when applied to cells during senescence induction (van Vliet et al., 2021), SISR represents a wholly novel phenotype of senescence since it occurs in established senescent cells exposed to acute and transient stressors. This observation has important implications for aging, where senescent cells are long lived and may survive multiple rounds of physiological stress, each of which could elicit SISR. Further investigation of SISR may thus reveal a previously unappreciated synergy between stress and senescence that exacerbates the already substantial inflammatory potential of senescent cells.

## Methods

### Cell culture

IMR-90 human lung fibroblasts (Coriell Institute) were cultured in high-glucose DMEM (Gibco) supplemented with 10% heat-inactivated FBS (Gibco), nonessential amino acids (1x, Gibco), and antibiotic/antimycotic mix (1x, Gibco). Cells at ∼40-60% confluency and at a population doubling level (PDL) between 20-30 were considered Cycling (Cyc). Contact-Inhibited Quiescent cells (Qui) were prepared essentially as described (Mitra et al., 2018) by allowing Cyc cells to reach ∼100% confluency and growing for an additional four days, changing media every 2 days and 24 hours before harvest. Etoposide-induced senescent (Etop) cells were prepared by treating Cyc cells with 50 µM etoposide (Selleckchem, dissolved in DMSO) over 10 days (treatments on day 0, day 2, and day 4) with media changed every two days. Senescence induction in WI-38 lung fibroblasts was conducted identically, while 25 µM etoposide was used for BJ human foreskin fibroblasts. Human Vein Endothelial Cells (HUVEC) were treated once with 10 µM etoposide and then incubated for 3 days before harvest. Ionizing-radiation induced senescent (IR) cells were generated as described (Neri et al., 2021) by exposing Cyc cells to 15 grays of ionizing radiation from a cesium-137 irradiator (Nordion) and then culturing for 10 days with media changed every two days. Serum-induced quiescence (SIQ) cells were generated by incubating cells in media as described except serum (FBS) was reduced to 0.2%; cells were cultured for at least four days in these conditions to establish quiescence with media changed every 2 days. Cells were harvested by trypsinization using TrypLE (ThermoFisher Scientific), pelleted at 4⁰C for 5 minutes at 600 x g and washed once in PBS before generation of protein lysates or bulk RNA as described below.

### Senescence validation

Senescence was validated using at least one of four methods: SA-β-Gal staining as per manufacturer’s instructions (Cell Signaling Technology), determination of cell cycle arrest using an EdU incorporation assay (ThermoFisher Scientific) as per manufacturer’s instructions, measurement of senescence associated protein markers (increased p16^INK4A^ and p21^CIP1^, decreased LMNB1), and elevated SASP factors measured by RT-qPCR (*CXCL8*, *IL6*, *CCL2*).

### Tandem Mass Tag mass spectrometry preparation and data analysis

Cyc, Qui, and Etop cells prepared as described were harvested by generating cell pellets through trypsinization and resuspended in mass spec buffer (100 mM Tris, 150 mM NaCL, 4% SDS, 1% Triton X-114, 0.1M DTT, and 1X Halt Protease Cocktail (Sigma) followed by sonication for 2.5 minutes and denaturation at 95⁰C for 3 minutes. Lysates were submitted for processing and analysis to Poochon Scientific (Frederick,MD) for quantitative proteomics using trypsin digestion, TMT-16plex labeling, fractionation by reverse-phase UHPLC, and LC-MS/MS analysis. Processed mass spectrometry data are available in **Table S1**.

Data was analyzed and volcano plots were generated using R data analysis software. Mass spectrometry data was filtered for proteins assigned with “medium” or “high” confidence at detection, as well as for proteins that showed a coefficient of variation ≤ 0.15 among biological triplicates. After filtering, principal component analysis was performed using base R function with all 9 sample libraries as the input. Biologically relevant protein changes were arbitrarily assigned to a log_2_ fold-change (log2FC) of 0.2/-0.2 and p-values ≤ 0.05 by two-tailed student’s T-test (Etop vs. Cyc, or Etop vs. Qui). GSEA analysis was performed on proteins identified as having at least 0.2/-0.2 log2FC in both the Etop/Cyc and Etop/Qui comparisons using the ClusterProfiler package(Wu *et al*., 2021).

### Western blot and RT-qPCR analysis

Protein lysates for western blot analysis were extracted from cell pellets (≤ 2 x 10^6^ cells) using 100 uL lysis buffer (2% SDS 50 mM HEPES) followed by water bath sonication and denaturation at 95⁰C for 3 minutes. Lysates were quantified using a BCA protein assay kit (ThermoFisher Scientific) and then equal amounts of protein were loaded onto a 4-20% SDS-PAGE gel (Biorad) and then transferred to a nitrocellulose membrane (BioRad). Western blot analysis was performed using the iBind system (ThermoFisher Scientific) as described by the manufacturer and with antibodies and amounts listed in Table S4. Quantification of western blots was performed in Image Lab (BioRad). Antibodies and dilutions used in this study listed in **Table S4**.

Bulk-RNA free of genomic DNA for RT-qPCR analysis was extracted from cell pellets (≤ 2 x 10^6^ cells) using the RNeasy plus spin-column purification kit (Qiagen) and performed using the Qiacube liquid handler (Qiagen) as described by the manufacturer. Extracted RNA was then amplified using Maxima reverse transcriptase (ThermoFisher Scientific) and random hexamer primer. cDNA was then analyzed using two-step RT-qPCR using SYBR Green mix (Kappa biosystems) and analyzed using the 2^-ΔΔCT^ method. Primers used are listed in **Table S4**.

### Ribosome Sequencing Library Preparation

IMR90 cell prepared as described above for Cyc, Qui, IR, and Etop conditions were harvested by transfer to ice and washing once with ice-cold 0.1 mg/mL cycloheximide in PBS. After supernatant was removed completely, cells were lysed by adding 500 uL ice-cold polysome lysis buffer (20 mM Tris pH 7.5, 50 mM NaCl, 1 mM DTT, 50 mM KCl, 1% Triton x-100, 0.1 mg/mL cycloheximide, and 1x HALT protease phosphatase inhibitor cocktail (ThermoFisher Scientific)) dropwise. Lysates were scrapped and then triturated 3x through a 26g needle followed by centrifugation at 18k x g at 4⁰C for 10 min. RNA content was approximated by nanodrop OD 260 units and 50 ug was aliquoted for RNA-sequencing and Ribo-sequencing for each biological replicate.

Ribo-seq lysates were treated with 1 uL of RNase I (ThermoFisher Scientific) at 4⁰C for 40 minutes with shaking and reaction was stopped by addition of 200 U SUPERaseIN (ThermoFisher Scientific). Digested lysates were then overlaid on 900 uL of a 1M sucrose cushion in a 13 x 51 mm polycarbonate tube (Beckman Coulter), followed by ultracentrifugation at 70k RPM at 4⁰C for 2 h (Optima TLX ultracentrifuge, Beckman Coulter). Ribosome pellets were then resuspended in Trizol and extracted for RNA according to manufacturer’s instructions. Extracted RNA was then subjected to DNAse I treatment for 15 minutes at 37⁰C and then subjected to gel purification on a 15% polyacrylamide TBE-urea gel to isolate ribosome footprints. Isolated footprints were extracted from gel slices using extraction buffer (300 nM NaOAc pH 5.5, 1 mM EDTA, 0.25% SDS) overnight at room temperature followed by isopropanol precipitation. Precipitated footprints were depleted of rRNA using Low Input RiboMinus Eukaryote System v2 kit (ThermoFisher Scientific) and then subjected to end healing using T4 PNK (New England BioLabs). Resulting footprints were then used to generate sequencing libraries using the NEBNext Small RNA Library Prep Set for Illumina kit (New England BioLabs). Library quality was assessed using High Sensitivity DNA Chips with the Agilent 2100 Bioanalyzer System (Agilent Technologies).

RNA-seq lysates were extracted for bulk RNA in Trizol according to manufacturer’s instructions. RNA quality was assessed using a TapeStation analyzer (Agilent Technologies) and only RNA integrity numbers (RIN) greater than 9 were chose for further library preparation. Libraries were prepared with the TruSeq Stranded mRNA library preparation kit (Illumina).

Sequencing of Ribo-seq and RNA-seq libraries were performed in multiplex separately using S1 Flowcell for the NovaSeq 6000 sequencer (Illumina). Demultiplexing was performed using bcl2fastq software (Illumina).

### Ribosome Sequencing Analysis

FASTQ files for Ribo-seq reads were trimmed for adaptors (3’*AGATCGGAAGAGCACACGTCTGAACTCCAGTCAC*) using cutadapt (v4.0) and then filtered for rRNA reads by aligning against human18S, 28S, 5S, and 5.8S rRNA sequences using Bowtie2 v2.4.5 and collecting unaligned reads with the “–un” argument. FASTQ files for RNA-seq reads were also trimmed for adaptors (3’AGATCGGAAGAGCACACGTCTGAACTCCAGTCA) using cutadapt (v4.0).

Processed RNA-seq and Ribo-seq reads were then aligned to the human genome (hg38 release, Ensembl GTF v108) using STAR (v2.7.10b) with “—quantMode” option enabled. Resulting count matrix was then used for translation efficiency analysis as previously described (Chothani et al., 2019) using DEseq2 v1.36.0 and design argument as “∼Condition+SeqType+Condition:SeqType”, where Condition was cell state (Cyc, Qui, Etop, IR) and SeqType was Ribo-seq or RNA-seq reads. RPKM analysis was obtained starting from the STAR generated read count matrix and then normalizing each gene total reads sequenced obtained from de-multiplexed fastq files prior to alignment; library-normalized reads were then transformed into RPKM by dividing reads by the max annotated transcript length (kb) for each gene.

Metaplots were generated by dividing each genomic region of a transcript (5’ UTR, coding sequence, 3’UTR) into 20 bins. The 5’ ends of the RPFs were then assigned to each of the bins based on the corresponding location of their ends. The binned read counts were normalized based on transcript expression (total number of reads in transcript) and were averaged across all transcripts. The standard error of the mean for each bin was calculated across replicates.

### Treatment of cells with proteotoxic and metabolic stress

Starvation media (-Met -Cys) was prepared from high glucose DMEM lacking glutamine, methionine, and cysteine (Gibco) and then supplemented with 20% dialyzed FBS and glutamine to typical DMEM levels. Amino acid starvation was then performed by removing media from cultured cells, washing once in PBS, and then replacing with -Met -Cys media followed by incubation for 3h unless otherwise indicated. Proteotoxic stress was induced using thapsigargin (Tg, Sigma) supplemented into culture media described previously at 25 nM and incubated on cells for 3 h unless otherwise indicated. Cells for either stress condition were harvested after treatment and then prepared for bulk RNA or protein lysates as described above, or for direct-RNA nanopore sequencing as described below.

### Direct-RNA Nanopore sequencing

Total RNAs were extracted from cells grown in 150 mM plates using Trizol according to manufacturer’s instructions. RNA quality was measured using a Tapestation gel electrophoresis system (Agilent) and RNA integrity numbers greater than 8 were considered suitable for analysis. RNA was then prepared for sequencing essentially as described for TERA5-Seq (Ibrahim *et al*., 2021). Briefly, ∼40 µg of bulk RNA was subjected to poly(A) selection using Oligo d(T)25 magnetic beads (NEB) followed by on-bead 5’ ligation of a biotinylated REL5 linker sequence using T4 RNA ligase 1 (NEB) for 3 h at 37⁰C. 500 ng of REL5-ligated poly(A) RNAs were used for library preparation using the direct RNA sequencing kit (Nanopore Technologies). Libraries were quantified using a Qubit 1X dsDNA High Sensitivity assay kit (ThermoFisher Scientific) and then analyzed on a FLO-MIN106 flow cell on a Minion device (Nanopore Technologies).

### Processing and analysis of direct-RNA long-read sequencing

Direct RNA sequencing data were basecalled using Guppy (v3.4.5). By protocol design, some reads contain a 5’ linker (REL5) that marks the in-vivo 5’ end. Cutadapt (v2.8) was used to identify and remove adaptors ligated on the 5’ end of reads using parameters -g AATGATACGGCGACCACCGAGATCTACACTCTTTCCCTACACGACGCTCTTCCGATCT -- overlap 31 --minimum-length 25 --error-rate 0. All reads except those smaller than 25nts are retained from further processing. Information about reads containing the adaptor is also retained for targeted analysis of 5’ end degradation. All reads were subsequently aligned against ribosomal sequences using minimap2 (v2.17). Reads that matched ribosomal sequences were excluded from further analysis. Filtered reads were mapped on reference human genome build 38 (hg38) using minimap2 (v2.17) with parameters -a -x splice -k 12 -u b -p 1 --secondary=yes. They were also aligned against the human transcriptome (Ensembl v91) with parameters -a -x map-ont -k 12 -u f -p 1 --secondary=yes. Transcript count was quantified as the number of reads aligning on each transcript. Differential expression of transcripts/genes was performed with DESeq2(Love et al., 2014).

### Poly-A tail and transcript length analysis

Nanopolish polya was used to extract poly(A) tail lengths from raw nanopore signal(Workman et al., 2019). Only poly(A) tail lengths that passed the software quality control scores and were tagged as “PASS” were used in further analysis. Differential analysis for poly(A) tail and read lengths across conditions was performed with NanopLen package using the linear mixed model option for quantification of statistical significance (https://github.com/maragkakislab/nanoplen). In-house python scripts were used for data visualization.

### Transcriptome isoform analysis

Isoform analysis from direct RNA sequencing data was performed using FLAIR (Tang et al., 2020). All libraries were merged for isoform identification in order to create a common reference for all downstream comparisons. Differential isoform usage across conditions was then performed with diff_iso_usage.py and diffsplice_fishers_exact.py. Hits that were common in both replicate comparisons were selected as high confidence isoforms.

### Transfection of *in vitro* generated mRNA

cDNA for *in-vitro* generated mRNA constructs was obtained from expansion of an ATF4 Mammalian Gene Collection *Escherichia Coli* strain (Horizon Gene Editing, Clone ID: 3454473), while G. Extracted DNA was PCR amplified using primers listed in **Table S4** to generate *ATF4* WT or Mut constructs, purified (PCR Cleanup Kit, Qiagen) and mRNA was generated using mMESSAGe mMACHINE T7 Ultra Kit (Ambion) according to manufacturer’s instructions. 500 ng of generated mRNA was transfected into Cyc or Etop cells using 4 µl Lipofectamine MessengerMAX Reagent (ThermoFisher Scientific). Cells were incubated for 12 hours until harvest for bulk RNA and protein extraction.

### Protein synthesis assay

Protein synthesis was measured using the Click-iT HPG Alexa Fluor 488 Protein Synthesis Assay kit (ThermoFisher Scientific). Cells were labeled in media as described above supplemented with 50 µM homo-paraglycine (HPG, methionine analogue) for times indicated in experiments. After labeling, cells were fixed and analyzed as per manufacturer’s instructions using a BZ-X (Keyence) fluorescent microscope. Protein synthesis was quantified by extracting the total brightness measured in the blue channel (fluorescein) from areas positive in the DAPI channel.

### Polysome profiling

Lysates for polysome profiling were prepared by treating cells with 100 µg/mL cycloheximide (Sigma) in culture media for 10 minutes at 37⁰C. Cells were then washed with ice-cold PBS also containing cycloheximide and lysed on ice by adding Polysome Buffer ( 20 mM Tris pH 7.4, 150 mM NaCl, 5 mM MgCl_2_, 1 mM DTT, 1 % v/v Triton X-100, 1x protease inhibitors (ThermoFisher Scientific), 20 U/mL DNase I (Roche), 100 µg/mL cycloheximide) dropwise to cells and then scraping. Lysate was then triturated through a 26G needle (BD) and centrifuged for 10 minutes at 20,000 x g at 4⁰C. RNA content was approximated using UV absorbance on a Nanodrop machine and 20 µg of RNA was loaded onto 5%-45% sucrose gradient and analyzed on a UV/VIS fractionator (Brandell). Raw absorbance readings were analyzed and plotted in R and normalized UV absorbance was calculated by dividing each measurement over the total absorbance measured.

For normalization of polysome lysates to cell count, adherent cells were washed with PBS and lifted with TrypLE (ThermoFisher Scientific), both containing 100 µg/mL cycloheximide. Lifted cells were resuspended in ice-cold PBS with 100 µg/mL cycloheximide and counted using a Cellometer Auto 2000 cell counter (Nexcelom). Lysates were then generated using Polysome Extraction Buffer (PEB) to a concentration of 20 million cells/mL. For pre-T1 analysis, gradients were loaded with lysate equivalent to 10 million cells and analyzed as described above. For post-T1 analysis, lysates equivalent to 10 million cells were incubated with 1 uL of RNAse T1 (ThermoFisher Scientific) for 40 minutes at RT before reaction was quenched on ice with 200 U of SuperaseIN (ThermoFisher Scientific). Digested lysate was then analyzed on a sucrose gradient as described above.

### siRNA Transfection

siRNAs were obtained from Horizon Gene Editing as ON_TARGETplus pools of four siRNAs. Cells were plated at 0.3 x 10^6^ cells per well in high-glucose DMEM (Gibco) supplemented with 10% heat-inactivated FBS (Gibco) and nonessential amino acids (1x, Gibco). siRNA (45 nMols) and 9 μL of RNAiMax transfection reagent (ThermoFisher Scientific) were incubated in 300 μL of OptiMem media (ThermoFisher Scientific) and then added dropwise to cells. Cells were incubated with siRNA mixture for 7 h and then media was replaced. Twenty-four hours after changing the media, cells were induced to senescence using 50 μM etoposide or harvested as cycling cells. Cells were harvested for protein and RNA after 3 days as described above.

### Measurement of Secreted Protein

Cells were plated at 0.3 x10^6 cells per well in a 6-well dish and treated with DMSO or thapsigargin for 2h the next day. Media was changed, and cells were grown for 24 or 48h as indicated, with media changed every 24h. At harvest, 1 mL of media was collected, placed on ice, and pelleted of cells and debris by centrifugation for 1 min 12k x g at 4⁰C; supernatant was collected at stored at −20⁰C until use. Conditioned media was analyzed for secreted protein using a Bio-Plex 200 (BioRad) multiplex ELISA machine using analyte plates manufactured by Luminex.

## Data availability

Sequencing data have been deposited in the Sequence Read Archive (SRA); accession: GSE223762 and GSE227766. Raw mass spectrometry data are available at the MassIVE data repository; accession: MSV000091077.

## Acknowledgements

This research was supported by the Intramural Research Program of the National Institute on Aging, National Institutes of Health and funded by grant NIH ZIA AG000696 to MM.

## Author contributions

MJP performed wet-lab experiments with assistance from JM, RM, CA, CB, JF, YP, SM, LW, and KA. SD performed computational analysis on dRNA-seq data. MJP performed computational analysis on mass spectrometry data and ribosome sequencing data. MJP and MM interpreted the data assisted by AR, NB and MG. MJP and MM wrote the manuscript with feedback from all authors.

