## Supplemental Figures for "Senescence suppresses the integrated stress response and activates a stress-enhanced secretory phenotype"

### Supplementary Figures

##### Figure S1

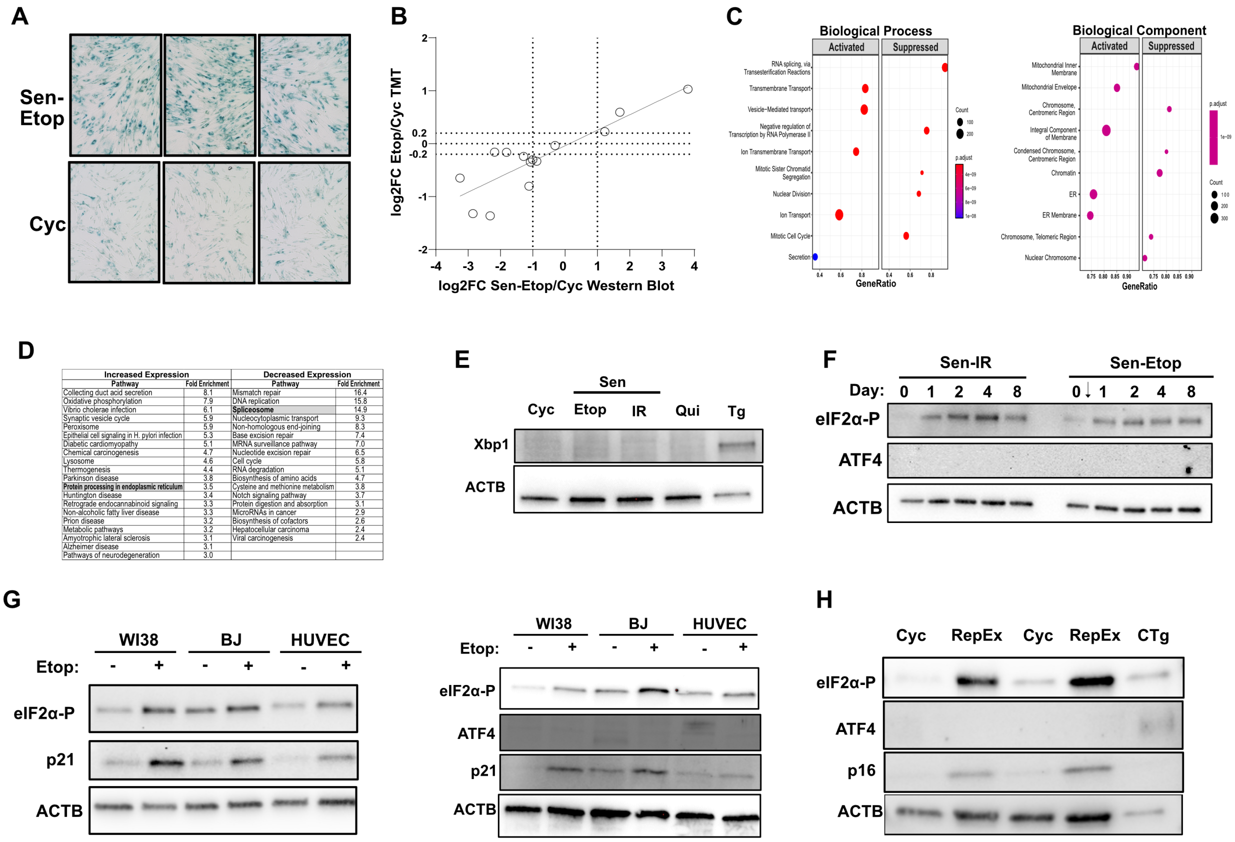


| Increased Expression | | Decreased Expression | |
|---|---|---|---|
| Pathway | Fold Enrichment | Pathway | Fold Enrichment |
| Collecting duct acid secretion | 8.1 | Mismatch repair | 16.4 |
| Oxidative phosphorylation | 7.9 | DNA replication | 15.8 |
| Vibrio cholerae infection | 6.1 | **Spliceosome** | 14.9 |
| Synaptic vesicle cycle | 5.9 | Nucleocytoplasmic transport | 9.3 |
| Peroxisome | 5.9 | Non-homologous end-joining | 8.3 |
| Epithelial cell signaling in H. pylori infection | 5.3 | Base excision repair | 7.4 |
| Diabetic cardiomyopathy | 5.1 | MRNA surveillance pathway | 7.0 |
| Chemical carcinogenesis | 4.7 | Nucleotide excision repair | 6.5 |
| Lysosome | 4.6 | Cell cycle | 5.8 |
| Thermogenesis | 4.4 | RNA degradation | 5.1 |
| Parkinson disease | 3.8 | Biosynthesis of amino acids | 4.7 |
| **Protein processing in endoplasmic reticulum** | 3.5 | Cysteine and methionine metabolism | 3.8 |
| Huntington disease | 3.4 | Notch signaling pathway | 3.7 |
| Retrograde endocannabinoid signaling | 3.3 | Protein digestion and absorption | 3.1 |
| Non-alcoholic fatty liver disease | 3.3 | MicroRNAs in cancer | 2.9 |
| Prion disease | 3.2 | Biosynthesis of cofactors | 2.6 |
| Metabolic pathways | 3.2 | Hepatocellular carcinoma | 2.4 |
| Amyotrophic lateral sclerosis | 3.1 | Viral carcinogenesis | 2.4 |
| Alzheimer disease | 3.1 | | |
| Pathways of neurodegeneration | 3.0 | | |

**Figure S1. (A)** Representative brightfield microscopy images of senescence-associated beta-galactosidase assay (n=6). (B) Correlation of log2FC measured by mass spectrometry and western blot analysis after normalization to ACTB for selected proteins (RPS20, p21, PCNA, LMNB1, eIF5, eIF2α, RPS10, eEF2, eEF2K, HSPA5, p16INK4A, RPL14, eEF1A, eIF4EBP1). Dashed lines on y-axis correspond to cutoff used to determine substantial change in mass spectrometry data, dashed lines on x-axis correspond to log2FC of 1/-1. **(C)** GSEA of proteins with uniquely increased and decreased expression in senescence. GSEA(Wu *et al.*, 2021) was performed for annotations in biological process (left) and cellular component (right) and p-value cutoff of 0.05. **(D)** KEGG analysis of proteins as above. **(E)** Western blot analysis for unfolded protein response (UPR) marker Xbp1 in cycling (Cyc), etoposide induced senescent (Etop), ionizing radiation induced senescent (IR), and Tg treated cells (Tg). **(F)** Western blot analysis of cells harvested at indicated days after treatment with IR or Etop. Arrow for Etop indicates day of treatment. **(G)** Western blot analysis of Etop induced senescence in three additional cell types: WI-38 lung fibroblasts (WI38, 50μM etoposide), BJ human foreskin fibroblasts (BJ, 25μM etoposide), and Human Vein Endothelial Cells (HUVEC, 10μM etoposide). Gels are technical duplicates. **(H)** Western blot analysis of replicative exhaustion induced senescent (RepEx) cells (PDL ~60).

Figure S2

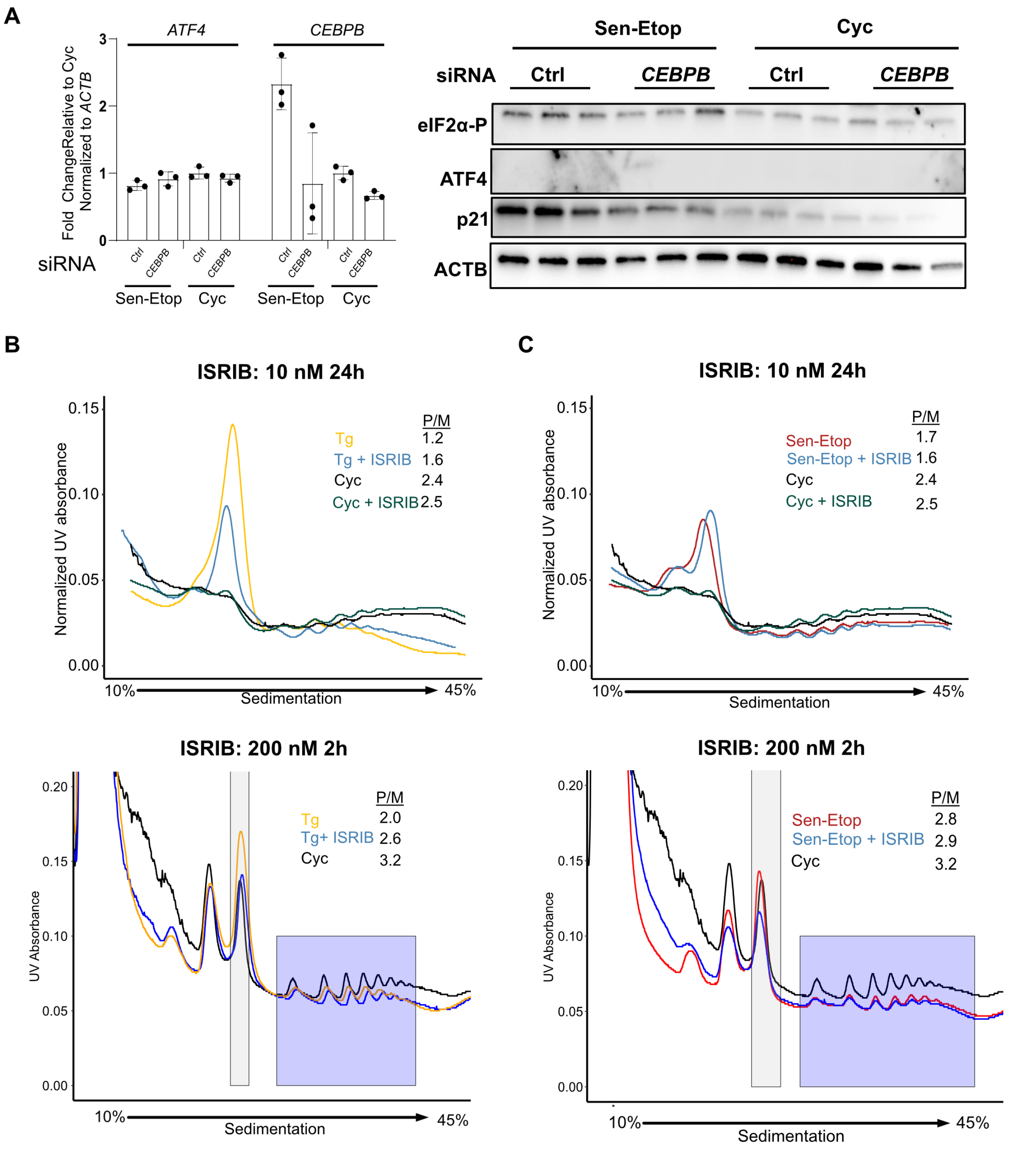


**Figure S2. (A)** At left, RT-qPCR of *ATF4* and *CEBPB* mRNA from etoposide-induced senescent (Sen-Etop) and cycling (Cyc) cells treated with siRNA against *CEBPB* or scrambled (ctrl). At right, western blot testing for ATF4 expression. **(B)** Polysome profiling of Cyc cells and cells treated with 25 nM thapsigargin for 3h (Tg); ISRIB applied at 10 nM for 24h (top) or 200 nM for 2h (bottom). **(C)** Polysome profiling of Cyc cells and Sen-Etop cells; ISRIB applied at 10 nM for 24h (top) or 200 nM for 2h (bottom).

#### Figure S3

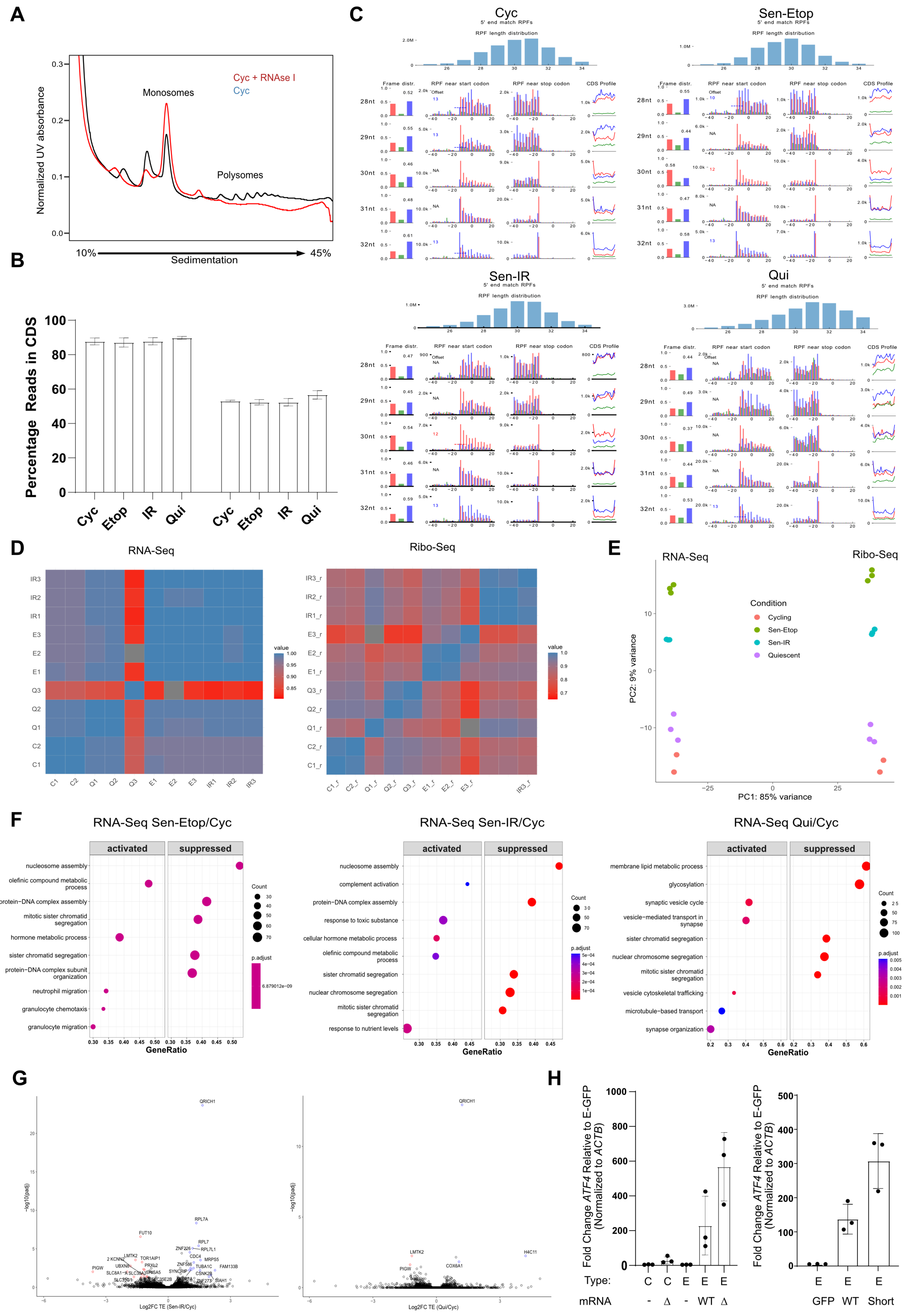

A
Normalized UV absorbance
Cyc + RNAse I
Cyc
Monosomes
Polysomes
10%
Sedimentation
45%
B
Percentage Reads in CDS
Cyc
Etop
IR
Qui
C
Cyc
Sen-Etop
Sen-IR
Qui
5' end match RPFs
RPF length distribution
Frame distr.
RPF near start codon
RPF near stop codon
CDS Profile
D
RNA-Seq
Ribo-Seq
value
E
RNA-Seq
Ribo-Seq
Condition
Cycling
Sen-Etop
Sen-IR
Quiescent
PC1: 85% variance
PC2: 9% variance
F
RNA-Seq Sen-Etop/Cyc
RNA-Seq Sen-IR/Cyc
RNA-Seq Qui/Cyc
activated
suppressed
Count
p.adjust
GeneRatio
nucleosome assembly
olefinic compound metabolic process
protein−DNA complex assembly
mitotic sister chromatid segregation
hormone metabolic process
sister chromatid segregation
protein−DNA complex subunit organization
neutrophil migration
granulocyte chemotaxis
granulocyte migration
complement activation
response to toxic substance
cellular hormone metabolic process
nuclear chromosome segregation
response to nutrient levels
membrane lipid metabolic process
glycosylation
synaptic vesicle cycle
vesicle−mediated transport in synapse
vesicle cytoskeletal trafficking
microtubule−based transport
synapse organization
G
QRICH1
Log2FC TE (Sen-IR/Cyc)
Log2FC TE (Qui/Cyc)
−log10(padj)
H
Fold Change ATF4 Relative to E-GFP (Normalized to ACTB)
Type:
mRNA
C
E
WT
Δ
GFP
Short

**Figure S3.** **(A)** Polysome profiling confirmation of ribosome footprinting; treatment of Cyc lysates with RNase I for 40 minutes at 4°C results in condensation of polysomes into monosomes without excessive degradation. **(B)** Percentage of reads in the coding sequence (CDS) for Ribo-seq and RNA-seq. **(C)** Ribo-seq quality control analysis indicating frame distribution (left column), read distribution at start and stop codons (center columns) and ratio of reads in the CDS for each frame (right column). Analysis performed using RIBOtish(Zhang et al., 2017) **(D)** Correlogram of RNA-seq and Ribo-seq libraries. **(E)** PCA of RNA-seq and Ribo-seq libraries; colors indicate cell type. **(F)** GSEA for biological process from RNA-seq data of indicated comparisons. **(G)** Volcano plot of differential translation efficiency for Sen-IR/Cyc (left) and Qui/Cyc (right) from ribosome sequencing. Statistically significant (adjusted p-value ≤ 0.05 by two-tailed t-test) log2FC values are colored as follows: log2FC ≤ -1, red; log2FC ≥ 1, blue; log2FC between -1 and 1, black. **(H)** RT-qPCR values for overexpression of *ATF4* mRNAs in Cyc (C) and Sen-Etop (E) cells; transfected mRNAs are none (-), WT ATF4 (WT), ΔuORF ATF4 (Δ) or Short-ATF4 (Short) as per Fig. 2G.

Figure S4

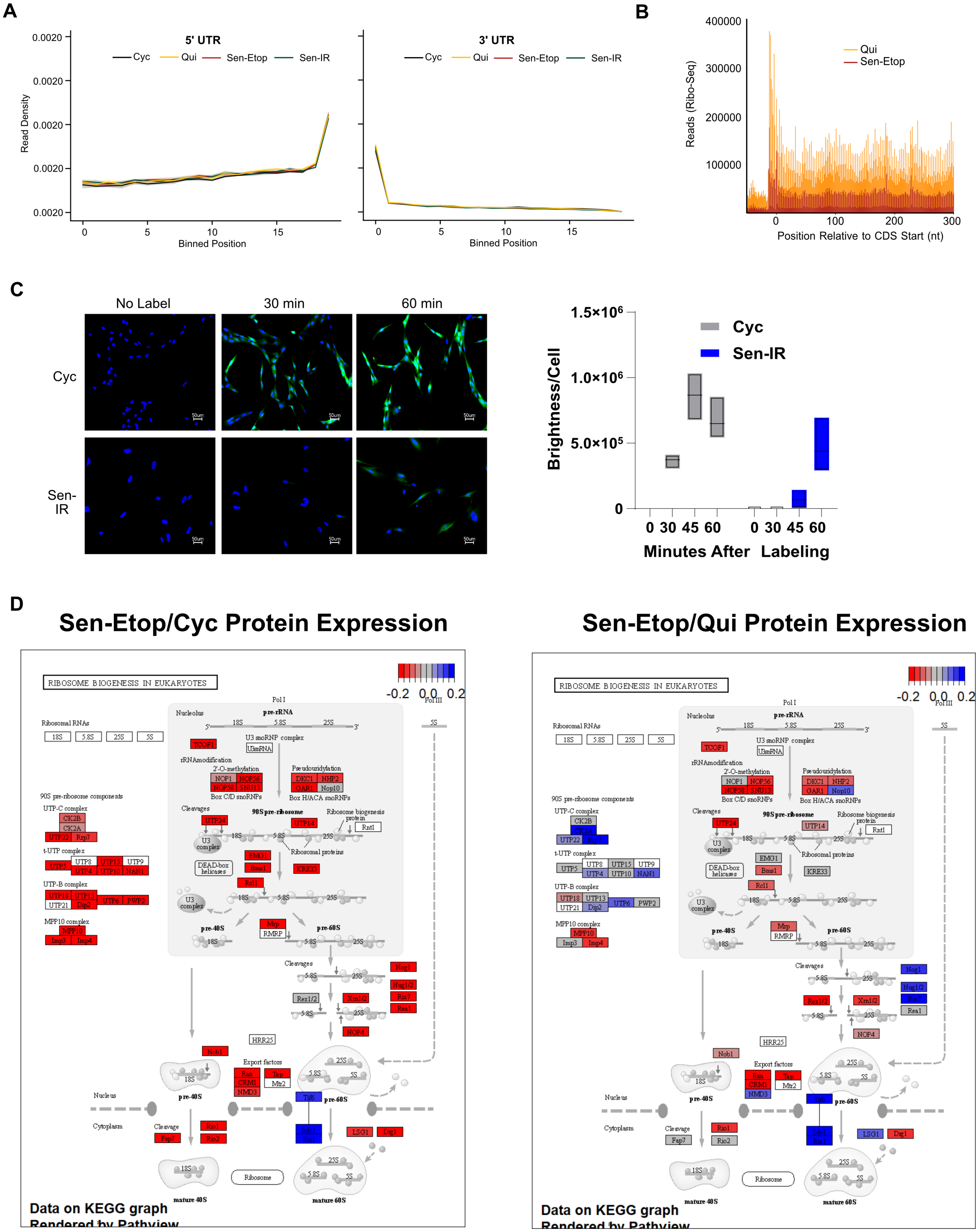
A
5' UTR
Cyc
Qui
Sen-Etop
Sen-IR
Read Density
Binned Position
3' UTR
B
Qui
Sen-Etop
Reads (Ribo-Seq)
Position Relative to CDS Start (nt)
C
No Label
30 min
60 min
Cyc
Sen-IR
Brightness/Cell
Minutes After Labeling
D
Sen-Etop/Cyc Protein Expression
Sen-Etop/Qui Protein Expression
RIBOSOME BIOGENESIS IN EUKARYOTES
Data on KEGG graph
Rendered by Pathview

**Figure S4. (A)** Metaplot analysis of ribosome footprints mapped to the transcriptome in the 5' and 3' untranslated regions (5' UTR and 3'UTR). Coordinates of footprints are aligned at the 5' end and then binned for a transcriptome-wide metalength (see methods). **(B)** Metaplot distribution of ribosome footprints near the CDS start site (-100 to +300). Reads from etoposide-induced senescent cells (Sen-Etop) contact-induced quiescent cells (Qui). **(C)** Representative images (n=4) from protein synthesis pulse-label assay (see methods) where time indicates length of pulse. Green, nascent protein; blue, DAPI stain. At right, quantification of translation as brightness detected within nuclear region divided by total DAPI count detected (Brightness/Cell) for ionizing radiation induced senescence (Sen-IR). **(D)** KEGG pathway analysis using Pathview for ribosome biogenesis of mass spectrometry data. Log2FC values derived from Sen/Cyc (left) and Sen/Qui (right) comparisons.

Figure S5

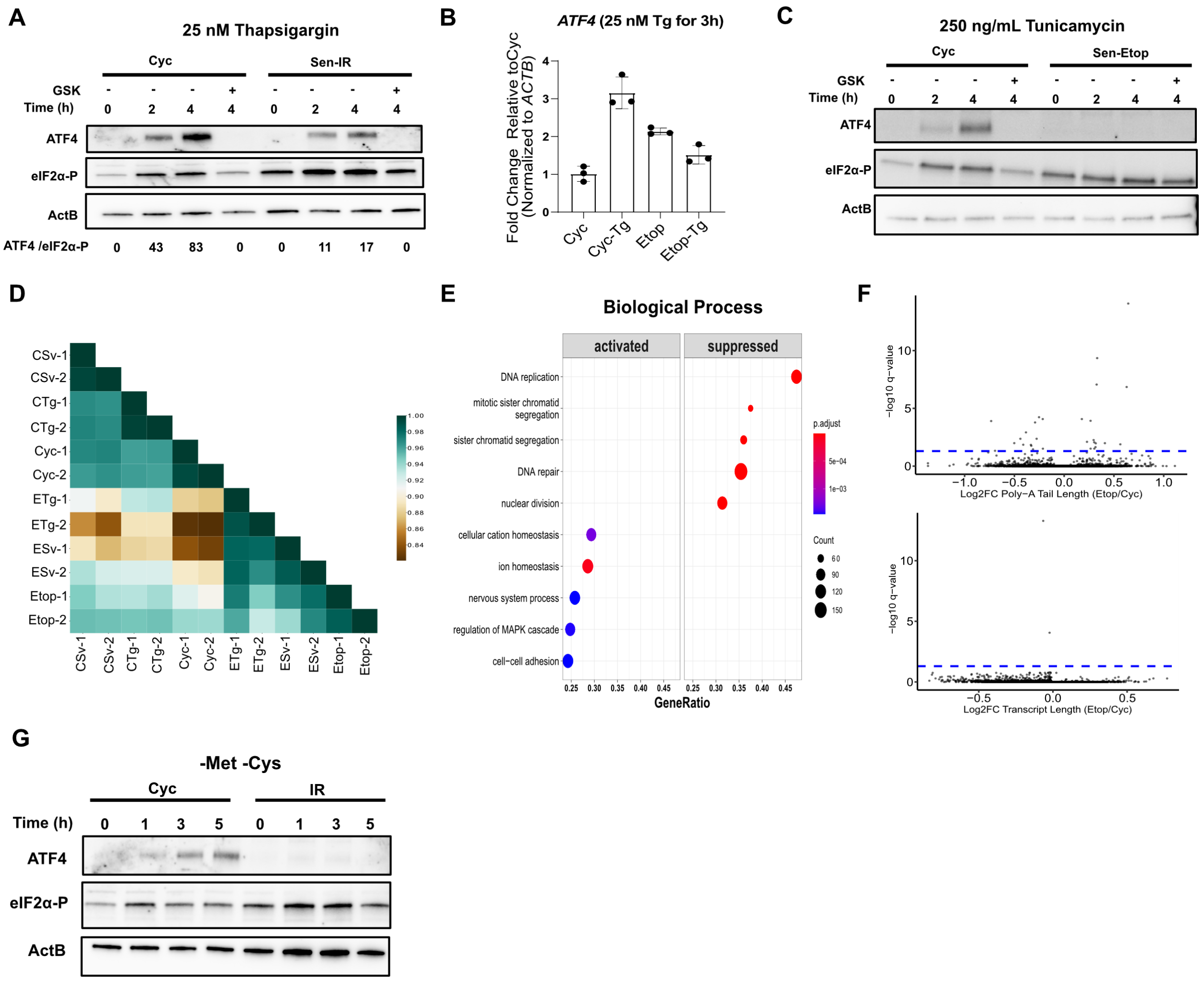


**Figure S5. (A)** Western blot analysis of cycling (Cyc) and ionizing radiation induced senescent (Sen-IR) cells treated with 25 nM Tg at indicated times. ATF4/eIF2α-P ratio calculated using densiometric analysis of shown western blots and rounded to nearest integer; GSK (550 nM) added concurrently with Tg for indicated lanes. **(B)** Western blot analysis of Cyc and etoposide induced senescent (Sen-Etop) cells treated with 250 ng/mL tunicamycin for indicated times; GSK applied as descrbed in (A). **(C)** RT-qPCR analysis for *ATF4* mRNA from bulk RNA extracted from Etop and Cyc cells with and without Tg treatment (3h, 25 nM). **(C)** Western blot of UPR markers cleaved ATF6 (ATF6c) and Xbp1. **(D)** Correlogram of dRNA-seq libraries. **(E)** GSEA for biological process of dRNA-seq Etop/Cyc comparison. **(F)** Volcano plots of log2 fold change for poly-A tail length (top) and transcript length (bottom) for Etop/Cyc dRNA-seq libraries obtained from NanopLen analysis (see methods). **(G)** Western blot analysis after starvation of Cyc and Sen-IR cells for methionine and cysteine amino acids cultured for indicated times.
